# Socioecological network adaptability enhances ecosystem service resilience in a tropical agricultural system

**DOI:** 10.1101/2025.09.05.674551

**Authors:** Anna Stanworth, Harry E.R. Shepherd, Kiole Imale, Kelvin S-H Peh, Rebecca J Morris

## Abstract

Understanding the vulnerability of ecosystem services to environmental change and species loss is critical to supporting communities dependent on services for well-being and livelihoods. However, assessments of ecosystem service resilience often overlook the complexity of socioecological systems, including the cascading effects of species loss and adaptive responses of biodiversity and humans.

We constructed a socioecological network representing ecosystem service provisioning in a tropical farming community, linking frugivore – plant interactions with provision of local services such as food and firewood, and simulated species loss under environmental change scenarios. We quantified the robustness of ecosystem services and identified key species and interaction pathways for such, and incorporated interaction rewiring into our simulations to assess the resilience of network and ecosystem services as species and humans adapt to environmental change.

Ecosystem service provisioning in our network was generally robust to simulated species loss. On average, our network could withstand the loss of over 80% of species before collapsing, with the loss of all services occurring in only the most extreme environmental change scenarios. When service loss did occur, such as the loss of firewood, it was driven by secondary extinctions after loss of primary species propagated through the network.

Interaction rewiring enhances ecosystem service resilience, allowing the loss of 18% more species before network collapse. Rewiring also altered the relative importance of individual species and their interaction pathways in maintaining resilient services. Species such as sago and coconut do not directly provide firewood and initially did not play an important role in maintaining robust firewood services with limited network connectivity. However, with rewiring these species helped maintain resilient firewood services in 20% more simulations, reflecting the importance of indirect interaction pathways and adaptivity.

These findings demonstrate that 1) species can maintain resilient ecosystem services even when they do not directly provide the service and 2) the adaptive capacity of humans and biodiversity can substantially enhance ecosystem service resilience. Overall, our study provides a framework for more accurately assessing the consequences of species loss on ecosystem services and human livelihoods by explicitly accounting for socioecological complexity and adaptability.

## 1. Introduction

Ecosystems and their services to people are under increasing pressure from climate change, habitat destruction, and species loss, which together threaten human livelihoods and wellbeing (Balvanera et al., 2019; IPBES, 2019). Predicting the impact of global change on ecosystem service provisioning is critical to understanding how human wellbeing and livelihoods will be impacted by future disruption (Bateman et al., 2013; Díaz et al., 2018; IPBES, 2019; Mouquet et al., 2015). However, there remains a limited understanding of how and when ecosystem services collapse (i.e., no longer supplied or declined to levels no longer of use to people) and their vulnerability to such failures (Keyes et al., 2021; Ross et al., 2021). Recognising the vulnerability of ecosystem services is essential for prioritising management actions that support communities at risk from species loss, particularly through the protection and restoration of species and habitats that underpin services such as food, medicine, and raw materials (Chaplin-Kramer et al., 2019; IPBES, 2019; Strassburg et al., 2020).

Ecosystem services are sustained by complex ecological and socioecological interactions (Stanworth et al., 2024). Species can support service provisioning both directly (e.g. pollinating crop species); and indirectly through their interactions with direct service providers or other supporting species (Keyes *et al*. 2021; Vitali *et al*. 2025). This ecological complexity makes it difficult to predict how species loss might affect ecosystem service delivery. For example, wild plant communities can support specialised predators that suppress agricultural pests, thus enhancing the pest control service for nearby crops (Tschumi et al., 2016). The loss of specific wild plant assemblages can then reduce natural pest regulation by diminishing predator populations, lowering food production. More broadly, the dense network of direct and indirect species interactions characteristic of natural systems means that species loss can trigger cascading impacts through secondary extinctions, where species with no direct link to the lost taxa can decline due to shared interaction partners (Ávila-Thieme et al., 2023; Dunne et al., 2002; Eklöf et al., 2013). These cascading effects on ecosystem services are generally underexplored (Domínguez-García & Kéfi, 2024; Keyes et al., 2021; Pocock et al., 2012), limiting our ability to predict how ecosystems may respond to species loss.

However, species interactions are not static. As species are lost, ecological niches open, promoting a shift in interaction partners and interaction strengths (CaraDonna et al., 2017; Marjakangas et al., 2025). Many species can adjust their interaction patterns, a process often referred to as interaction rewiring (Costa *et al*. 2018; Kaiser-Bunbury *et al*. 2010). The ability for interactions to rewire under changing conditions can buffer ecosystems against the functional consequences of species loss and enhance the robustness of service provisioning. For example, studies on common Robin (*Erithacus rubecula*) have shown that individuals increase dispersal frequency of alternative species after experimental loss of their main food source from a seed dispersal network, blackberries (Costa et al., 2018). This demonstrates that rewiring of interactions can confer resilience in communities as species adapt to shifting competition dynamics (Ives & Cardinale, 2004; Timóteo et al., 2016). However, the impact of interaction rewiring on ecosystem service robustness and resilience is also underexplored (Keyes et al., 2021). Crucially and less widely considered, rewiring can also occur within socioecological systems. Local communities adjust their interactions with biodiversity in response to environmental change, such as redistributing their dependence on different species through crop selection or harvesting different natural resources (Marjakangas et al., 2025). Adaptive responses of both taxa within networks and the sociological context in which services are provided could enhance the robustness and resilience of systems. Predicting the impact of species loss on ecosystem service provisioning therefore requires consideration of both secondary extinctions and interaction rewiring. However, many studies examining the impacts of species loss focus primarily on species directly providing services, without explicitly considering the secondary impacts on interaction pathways or rewiring of interactions, including human adaptive responses to environmental change Furthermore, ecosystem service studies often overlook the impact of human values, decisions, and adaptations that shape how people interact with biodiversity and influence service provisioning (Keyes et al., 2021; Ross et al., 2021).

Network ecology offers a framework for incorporating greater ecological complexity in assessments of how species losses may directly and indirectly affect ecosystem services (Stanworth et al., 2024). In particular, network-based approaches allow for the integration of ecological and socioecological processes into a unified socioecological network, with explicit measures describing how different components are connected, for example, through biodiversity-mediated interactions. This integration of processes allows researchers to conceptualise and quantify ecosystem service flows from biodiversity to people and reveal both direct and indirect interaction pathways that support ecosystem services (Stanworth et al., 2026). Network ecology can also allow the incorporation of interaction rewiring into assessments of ecosystem service provision. By modelling how species and people adjust their interactions, ecologists can infer the resilience of ecosystem services after a system has adapted to change, with resilience defined as the maintenance of service provisioning despite turnover in interactions (Marjakangas et al., 2025). By incorporating interaction rewiring into future scenario modelling, ecologists can identify species that should be targeted in conservation and management beyond those that are currently threatened or directly providing services, as well as those whose loss would disproportionately reduce service robustness (Rey et al., 2024). Moreover, by including information on the replaceability of service providers within networks, it is also possible to make more nuanced predictions about both the robustness and resilience of ecosystem services. A network approach can explicitly account for humans’ adaptive capacity to change and the potential reorganisation of socioecological interactions (Marjakangas et al., 2025; Vizentin-Bugoni et al., 2020). Thus, network ecology provides a valuable avenue for incorporating ecological complexity and adaptability into assessments of ecosystem service robustness.

Here, our objective is to show how modelling species loss within the network framework can reveal insights into the effects of species loss propagating through direct and indirect interaction pathways to determine ecosystem service resilience (Stanworth et al., 2026; Fig. 1). To do this, we construct a socioecological network representing a tropical agricultural system that connects frugivore-plant interactions and socioecological interactions between plants and local ecosystem service provisioning. We then apply ‘knockout extinction models’ to simulate primary extinction of species within the network (Bane et al., 2018) and estimate the robustness of individual ecosystem services to biodiversity loss driven by climate change, habitat degradation, and disease – three main components of global anthropogenic change (IPBES, 2019; Mahon et al., 2024). Our modelling framework incorporates the potential for both ecological and socioecological interaction rewiring, allowing us to infer ecosystem service resilience under more realistic scenarios of species loss. In doing this, we address three key objectives:

1. Quantify the robustness and resilience of individual ecosystem services to species loss.
2. Identify key species and interaction pathways that maintain robust and resilient ecosystem services and assess how these roles relate to direct service provisioning.
3. Assess how interaction rewiring influences the robustness and resilience of ecosystem services.

**Figure 1:**
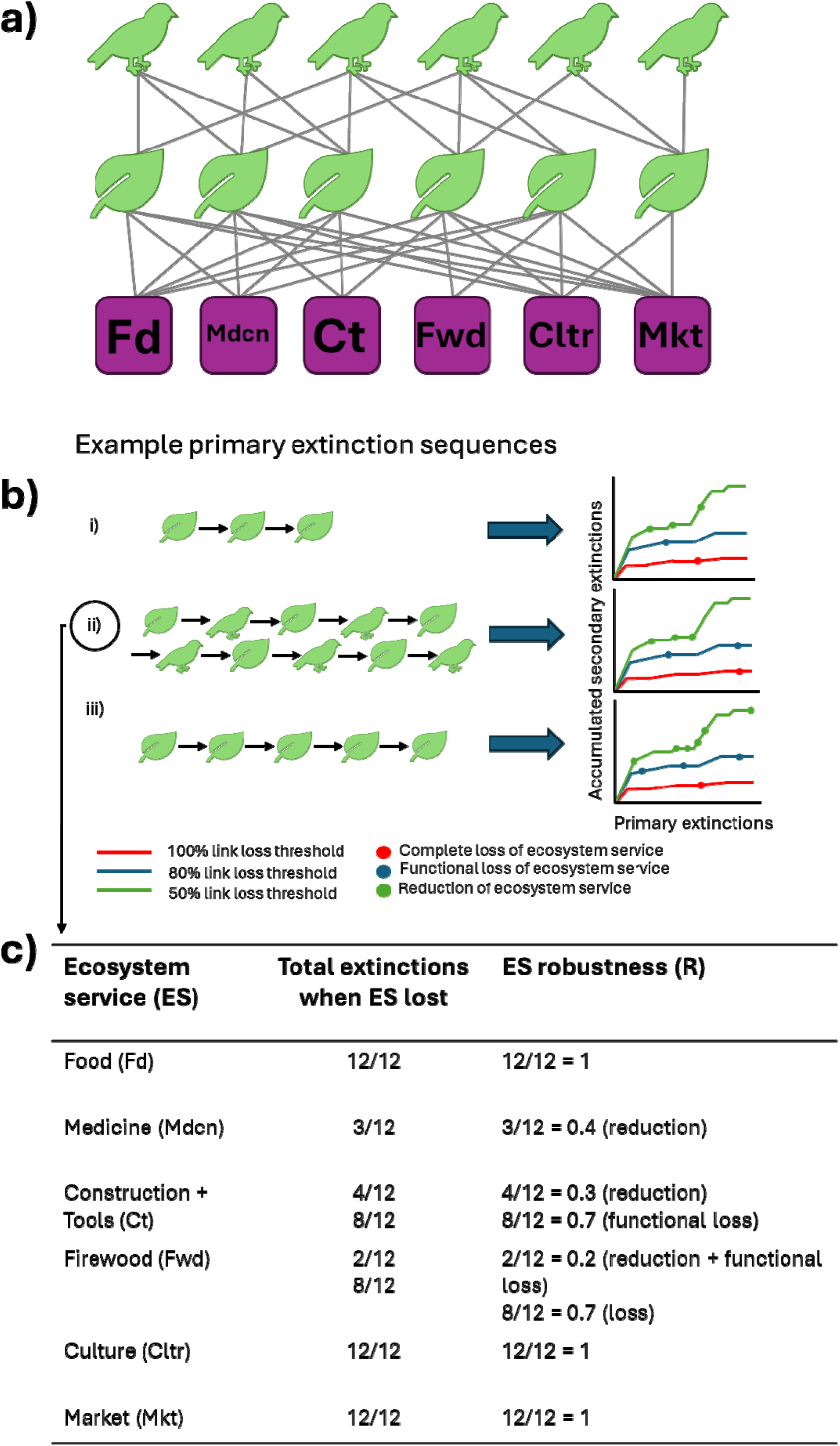
Conceptual representation of the extinction simulations and subsequent quantification of ecosystem service robustness. (a) The tripartite network represents frugivorous interactions between frugivores and plants, and the provision of ecosystem services between plants and the specific ecosystem services that they provide to the local community, including food (fd), medicine (mdcn), construction and tools (Ct), firewood (fwd), culture (cltr), and income from services (mkt). (b) This tripartite network undergoes various species loss simulations that result in primary and secondary extinctions of species and interaction pathways, and secondary extinctions of ecosystem services after different levels of link loss. Here, we show how different extinction sequences (i, ii, iii) translate to accumulated secondary extinction graphs. (c) The robustness of each ecosystem service is then calculated using the species loss that the service can withstand (i.e. accumulated primary and secondary extinctions) before it is lost, functionally lost, or reduced. Market refers to the market-mediated provision of services.

Taken together, this approach advances on previous assessments of ecosystem service resilience by explicitly integrating ecological complexity, sociological dynamics, and adaptive interaction rewiring. It demonstrates how individual ecosystem services and key species and interactions underpinning them can be identified within a realistic, socioecological system.

## 2. Methods

### 2.1 Dataset

We use data collected in the smallholder farming village of Ohu, Papau New Guinea (PNG) (S 05°13.081’, E 145°40.735’). Ohu is predominantly a farming community focused on subsistence food gardens and cash crop blocks, as well as harvesting wild plants from surrounding forests to complement nutritional requirements; and meet medicinal, construction, firewood and cultural needs (Bourke & Harwood, 2009; Hazenbosch et al., 2021). Data on observed ecological interactions and farmer – plant socioecological interactions was collected using household interviews, with 40 households interviewed by PNG staff from the New Guinea Binatang Research Centre (NGBRC) (Stanworth et al., 2026). In summary, each household was asked questions about 1) Socioecological demands (for example, plant species that provide services directly through cultivation and harvest); and 2) Species’ interactions that have been observed in the surrounding environment, with a focus on plant-frugivore interactions. All interviews were conducted in Tok Pisin or Amele (the local language in Ohu). All participants gave their prior-informed consent through verbal consent and were given the same opportunities to propose observed interactions. The interview responses were translated into English by NGBRC and the interview protocol was approved by the University of Southampton Research Ethics Committee under permit number 78769.A1. Details on the complete data collection, including specific questions provided to the interviewees, can be found in Stanworth et al., (2026).

### 2.2 Socioecological network

We constructed a tripartite-structured network composed of two layers of interactions, using the answers provided by participants as described in the previous section. The first layer depicts plant-frugivore interactions and was devised by linking observed interactions described by participants between plants (both utilised and non-utilised for service provisioning) and avian and bat frugivores. The second layer depicts plant-ecosystem service interactions and was devised by linking useful plants to the services they provide to participants in the local community (Fig. 1a). The plant-ecosystem service interaction layer was devised by linking useful plants to the services they provide to the local community, categorised into six ecosystem services: food, medicine, construction and tools, firewood, culture, and income from services.

Plants in the income from services category are sold at local markets and generate income through their market-mediated provision of different services. As plants can provide multiple services, species were not restricted to a single category.

Interactions in both layers are undirected, meaning that the observed interactions are assumed to be mutualistic (Stanworth et al., 2026). Plant-frugivore interactions were weighted by the regularity with which interactions were reported in interviews and, where applicable, published literature (Ong et al., 2021; Stanworth et al.2026). Plant-ecosystem service interactions were weighted by the regularity with which a service was attributed to a plant in interviews. We used the R package ‘network’ (Butts, 2008) to construct our tripartite network from an edge list of interactions between three sets of nodes: frugivores, plants, and ecosystem services incorporated directly in the network as nodes (Felipe-Lucia et al., 2022; Keyes et al., 2021).

### 2.3 Extinction scenarios and simulations

We created extinction sequences to simulate species loss under climate change, habitat loss, and crop diseases. To do this, we developed three scenarios, split into five sub-scenarios (Table 1), that represent realistic predictions of environmental change at our study site and the potential for cumulative impacts of global change on local biodiversity. Scenario 1 simulates climate change-driven declines in plant diversity of either 80% or 60% reductions in species richness. This reflects expected losses under the RCP 8.5 and 2.6 emission scenarios for the lowland rain and freshwater swamp forest ecoregion in Papau New Guinea, where Ohu village is located (Cámara-Leret, Raes, et al., 2019). Scenario 2 reflects biodiversity loss across multiple trophic levels (e.g., plants and frugivores) associated with human populations and increased forest encroachment (Bourke & Allen, 2021; Hazenbosch et al., 2022). Scenario 3 combines the effects of climate change, disease, and overharvesting to simulate the compounding pressures on local biodiversity. In this scenario, the species loss percentages in Scenario 1 are paired with the definite loss of *Cocos nucifera* (coconut), *Musa sp.* (banana), *Areca catechu* (betelnut), and *Colocasia esculenta* (taro) due to the spread of phytoplasma-associated diseases such as Bogia coconut syndrome (Gurr et al., 2016; Hazenbosch et al., 2022), and the loss of *Metroxylon sagu* (sago) due to over-harvesting (Hazenbosch et al., 2022).

**Table 1:**
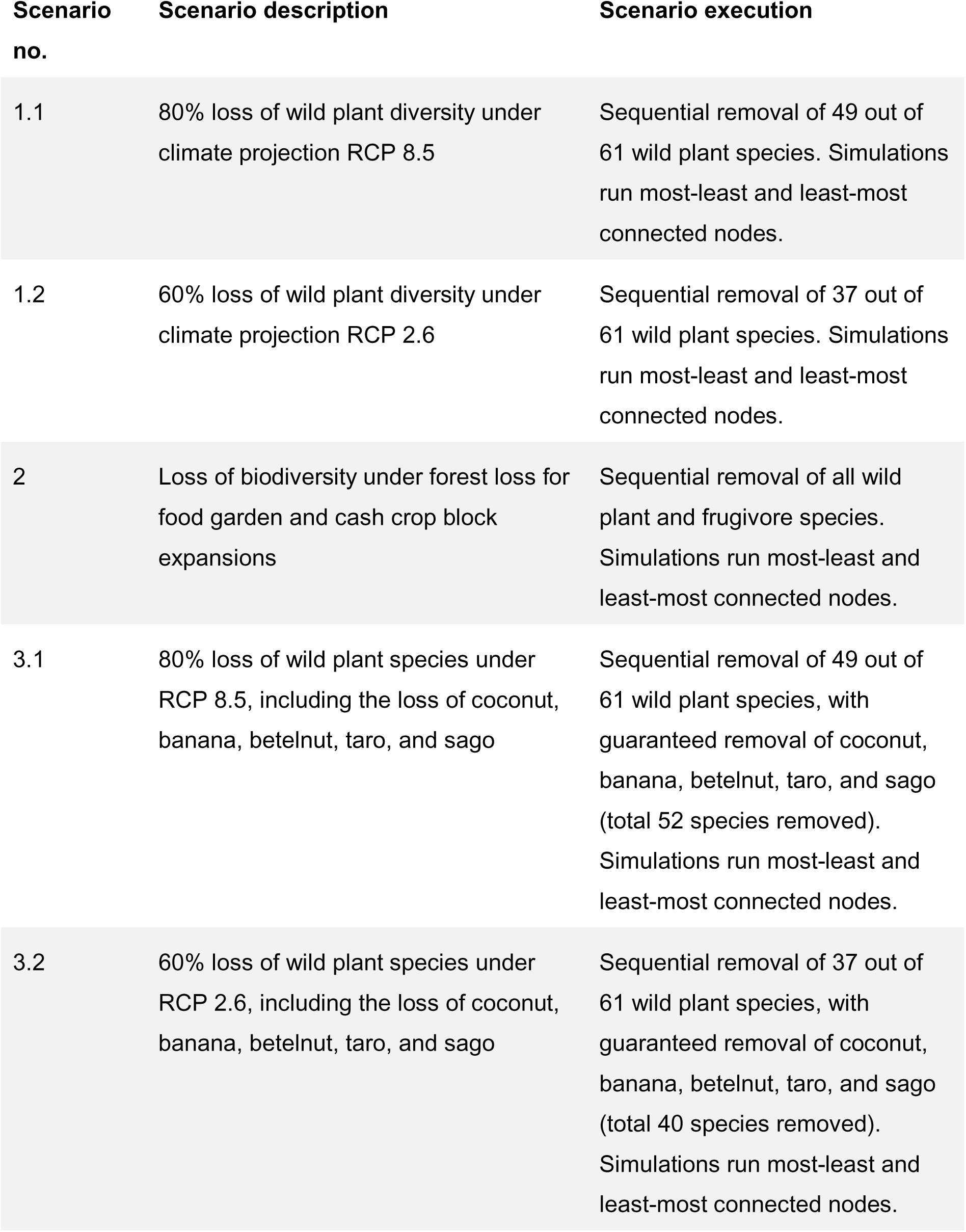
Description of five future change scenarios. For further details, see Table S1.

Extinction scenarios were simulated using the R package ‘NetworkExtinction’ (Ávila-Thieme et al., 2023). Each scenario began with the primary removal of a species from our socioecological network (i.e., either plant or frugivore depending on the scenario; Table 1). After each primary node was removed, we recorded all the resulting secondary extinctions to capture how species loss propagated through the network (Fig. 1b). We then removed the next species from the extinction sequence and again documented any secondary extinctions, repeating the process until the designated extinction thresholds for each scenario had been reached (Table 1). The primary extinctions orders were determined by the connectivity of each species within the network, measured using the degree of the node and calculated using the R package ‘igraph’ (Csárdi et al., 2025). Species were removed either (i) from most-to-least connected; or (ii) from least-to-most connected, reflecting the worst- and best-case scenarios of species loss, where species expected to have the most cascading impacts within a network (i.e., highest node degree) either lost first or last (Memmott et al., 2004). To account for ties in the connectivity, each extinction order was randomly shuffled 100 times. We then used the extinction orders with the highest number of secondary extinctions and lowest network robustness to run ecosystem service robustness and key species analysis. This represents the maximum impact our simulations had on network structures and the most conservative estimate of global change impacts on our socioecological network.

Each extinction scenario and primary extinction order were simulated three times, using secondary extinction thresholds of 50%, 80%, and 100% link loss to represent different levels of ecosystem service decline (Fig. 1b) (Ávila-Thieme et al., 2023; Jones et al., 2023).

Specifically, a node is lost from the network after losing 50%, 80% or 100% of its links, depending on the link loss threshold set. These thresholds reflect that disturbance is often gradual, where biodiversity is lost incrementally over time, for example, through forest encroachment (Sodhi et al., 2010; Tilman et al., 1994). By applying multiple thresholds, we capture how ecosystem services respond at different stages of biodiversity loss: from an initial reduction of services as biodiversity begins to decline (50% threshold), the functional loss of services as biodiversity falls below levels required to sustain them (80% threshold), and the complete collapse of services when all species directly supporting ecosystem service have been lost (100%) (Ávila-Thieme et al., 2023; Jones et al., 2023). This combination of three extinction scenarios, with five sub-scenarios and run most-to-least and least-to-most at three different link loss thresholds, resulted in 30 unique individual simulations (Table S1).

Each of the 30 unique simulations was then repeated twice: once allowing interaction rewiring, reflecting adaptive responses of both frugivores and farmers to species loss (Ávila-Thieme *et al*. 2023; Borah & Beckman 2024; Kaiser-Bunbury *et al*. 2010); and once with fixed interactions where no adaption was permitted, resulting in 60 total simulations. To do this, we incorporated interaction strength rewiring into our simulations (Ávila-Thieme et al., 2023; Marjakangas et al., 2025). When rewiring is incorporated and a node is lost from the network, its interactions are redistributed among remaining links according to its rewiring probability (Ávila-Thieme et al., 2023), reflecting ecological processes observed in plant–pollinator (DomínguezlZlGarcia et al., 2026) and seedlZldispersal networks (Costa et al., 2018). For example, if a plant providing 10% of food provisioning was lost from the network, that 10% of food provisioning would be redistributed to other plants, depending on the probability that other plants provide food services. For frugivore-plant interactions, rewiring probabilities were estimated using latent-trait modelling in the R package ‘cassandRa’ which infers latent interaction probabilities from the observed network structure (Terry, 2019). For plant-ecosystem service interactions, rewiring probabilities were derived from the likelihood that each plant species provides particular services to the local community, based on plant provisioning scores obtained from interviews with local farmers in Ohu (Stanworth et al., 2026). This meant that the rewiring probabilities for sociological interactions are grounded in local ecological knowledge and community preferences and priorities for crop and wild plant uses.

### 2.4 Data analysis

#### 2.4.1 Network and ecosystem service robustness analysis

To quantify the overall robustness of the network to species loss, we extracted the *R100* value (Ávila-Thieme et al., 2023), which indicates the fraction of primary extinction events that cause total collapse of the network after both primary extinctions and subsequent secondary extinctions (Ávila-Thieme et al., 2023). Values closer to 1 would indicate that all species must be lost before the network collapses, and therefore a perfectly robust network. To quantify the robustness specifically for ecosystem service supply to species loss, we calculated the percentage of species an ecosystem service node could withstand to lose before 50%, 80% or 100% of its links were lost within the network and the ecosystem service went secondarily extinct (Fig. 1c) (Keyes et al., 2021; Ross et al., 2021). This robustness value, *R,* was calculated by dividing the number of species extinctions that had occurred at the point at which an ecosystem service node was lost by the total possible number of species that could be lost (n=117).

#### 2.4.2 Roles of species in determining the robustness and resilience of ecosystem services

To identify which species and interaction pathways underpin the robustness and resilience of ecosystem services to species loss, we identified those whose loss initiates cascading effects through the network. We considered a species valuable for maintaining robust ecosystem service provisioning when the service robustness decreased after a species was removed early in the extinction sequence, indicating that its removal can trigger the loss of direct and indirect partners through interaction pathways (Domínguez-García & Kéfi, 2024; Pocock et al., 2012).

Therefore, a species was classified as valuable when it was lost in the first half of primary and secondary extinctions within a simulation, and *R* < 0.5. To quantify this, we assign each species a relative importance score for maintaining the robustness and resilience of different ecosystem services. This score was calculated as the proportion of extinction scenarios (out of 30) in which a species was classified as valuable.

## 3. Results

### 3.1 Network and ecosystem service robustness

We first consider the robustness of our socioecological network without interaction rewiring, thereby evaluating the response of species and services within our network to species loss in the absence of adaptive changes. Our network showed consistently high robustness across extinction simulations (Fig. 3; Table 2). On average, the network could withstand the loss of approximately 80% of its species before collapsing (*R100* =0.82 ± 0.03 SE; Fig. 3a; Table 2). Only under a select few simulations did our network robustness (*R100*) go below 0.5 (Fig. 3a), whereby complete functional loss (i.e. network collapse) occurred after fewer than half of all species were lost. These results were only found in simulations in which environmental change exerted compounded impacts on the network (Scenario 3, Table 1). Under these conditions, all six ecosystem services were lost, but only when species loss followed worst-case scenarios and loss thresholds were at their most conservative (50%) (Fig. 4a; Table 2). Overall, this indicates that although species loss may reduce ecosystem service provisioning, ecological functioning in our network is largely resilient except under extreme environmental change scenarios.

**Figure 2:**
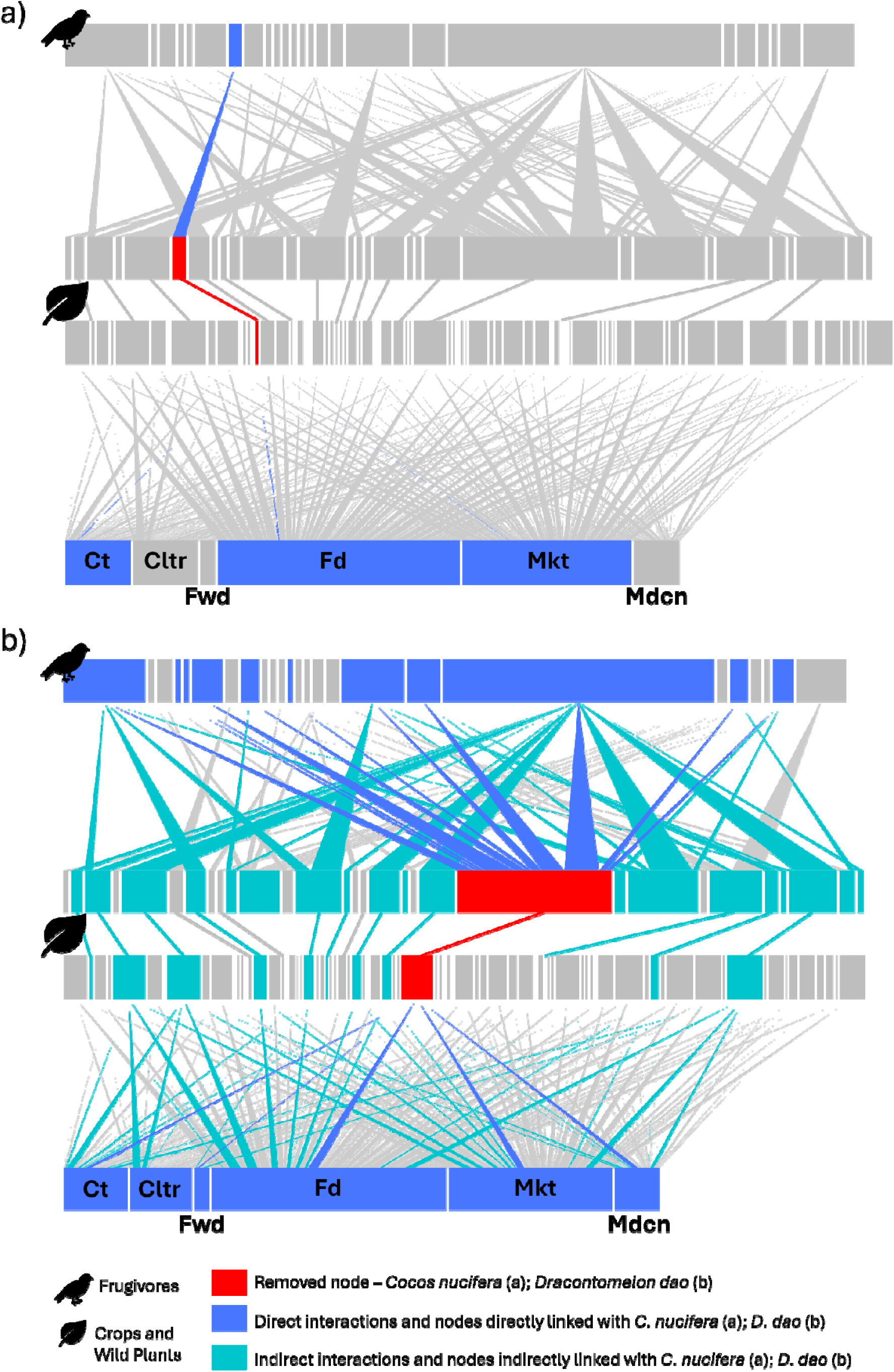
Socioecological tripartite networks illustrating direct (blue) and indirect (turquoise) interaction pathways following the loss of Cocos nucifera (a) and Dracontomelon dao (b), without interaction rewiring. The removal of C. nucifera has minimal impact, involving a single frugivorous interaction and limited direct ecosystem service provision (blue links are difficult to discern due to their thinness), with no secondary effects observed. In contrast, the loss of D. dao produces widespread effects throughout the network, impacting multiple direct and indirect pathways. As a resource for highly connected frugivores, its removal also affects other plant species that share these interaction partners, leading to cascading indirect impacts on overall ecosystem service provision.

**Figure 3:**
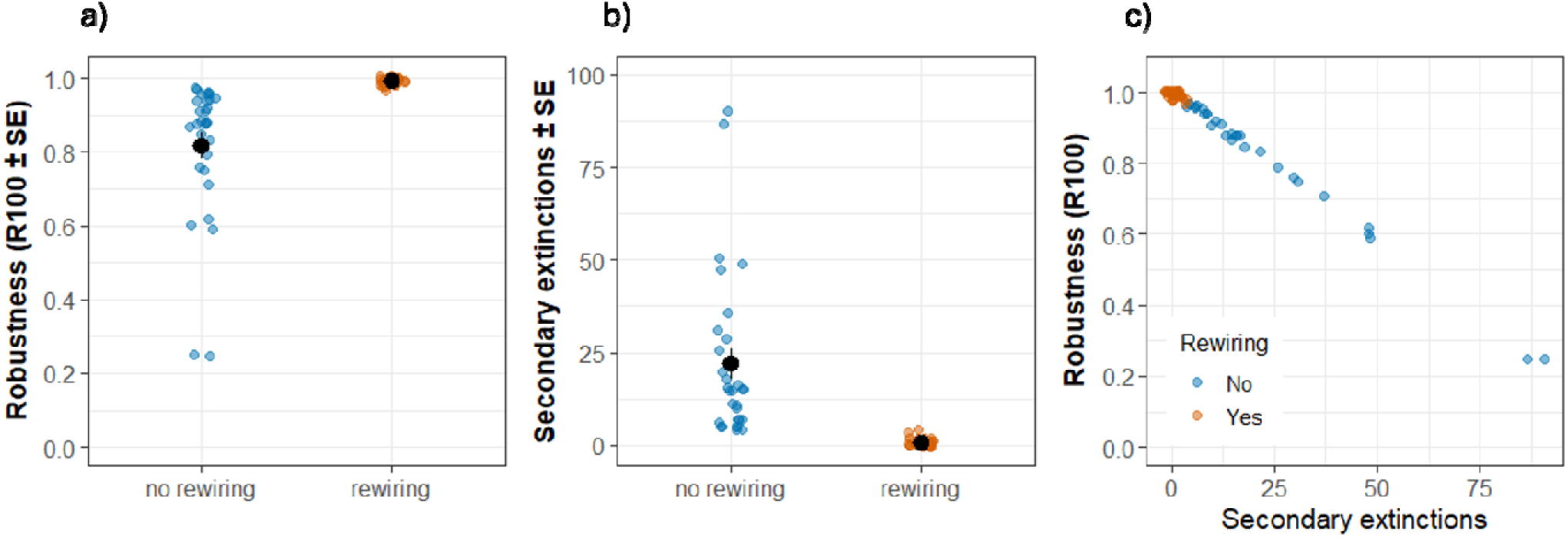
**Network robustness and secondary extinctions with and without interaction rewiring incorporated into extinction scenarios, and their relationship**. (a) Overall network robustness (R100); (b) number of secondary extinctions, both without interaction rewiring and with rewiring; and (c) the relationship between secondary extinctions and overall network robustness (R100). R100 refers to the percentage of primary extinction events that cause 100% of all species and ecosystem services in the network to go extinct after both intentional primary extinctions and subsequent secondary extinctions. Raw data points represent individual extinction scenarios (n = 30). Data points are slightly jittered to aid visualisation.

**Figure 4:**
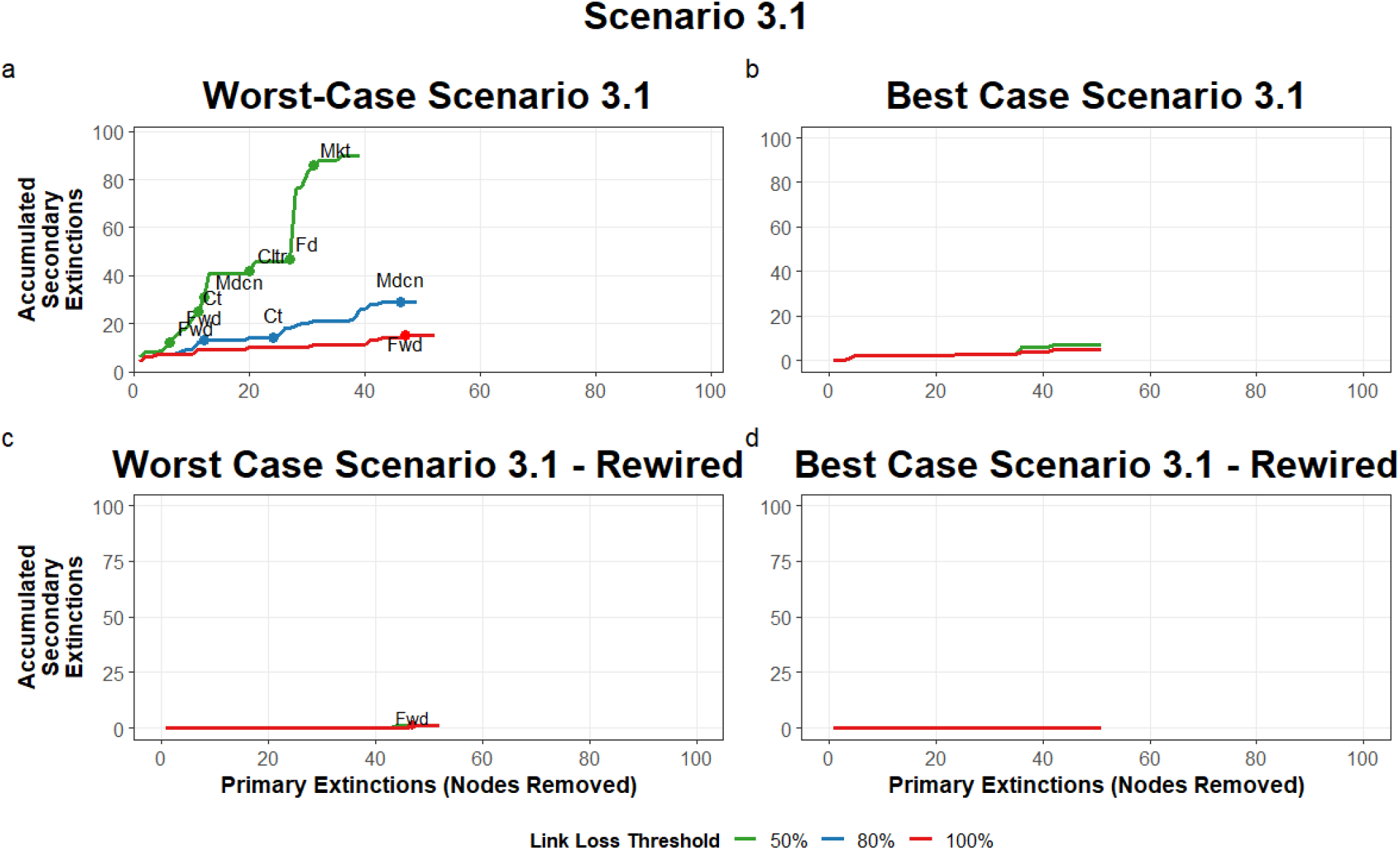
Accumulated secondary extinctions under an extreme environmental change scenario in which the primary extinction of up to 80% of wild plants and key harvested species is simulated. Simulations vary depending on whether interaction rewiring is excluded (top row) or incorporated (bottom row); and whether species extinctions follow worst-case (left column) or best-case scenarios (right column). Link-loss thresholds indicate the percentage of links a node (i.e. species or ecosystem service) can lose before being removed from the network Extinction points for each ecosystem service are labelled; with fd = food, mkt = income from services, mdcn = medicine, ct = construction and tools, cltr = culture, and fwd = firewood. In all panels except A, secondary extinctions follow similar trajectories across link-loss thresholds; therefore, lines are layered and not all thresholds are visible.

**Table 2.**
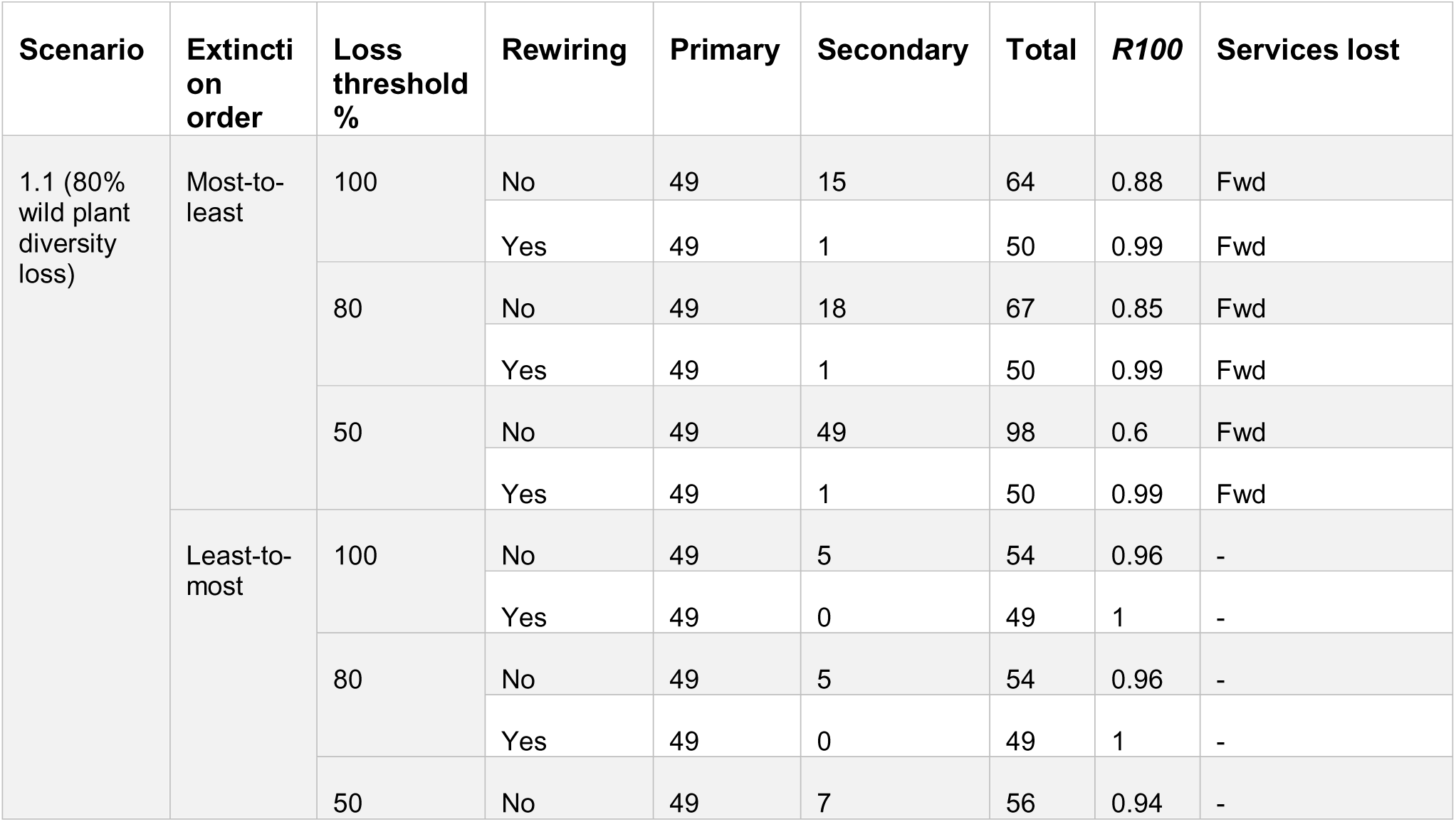
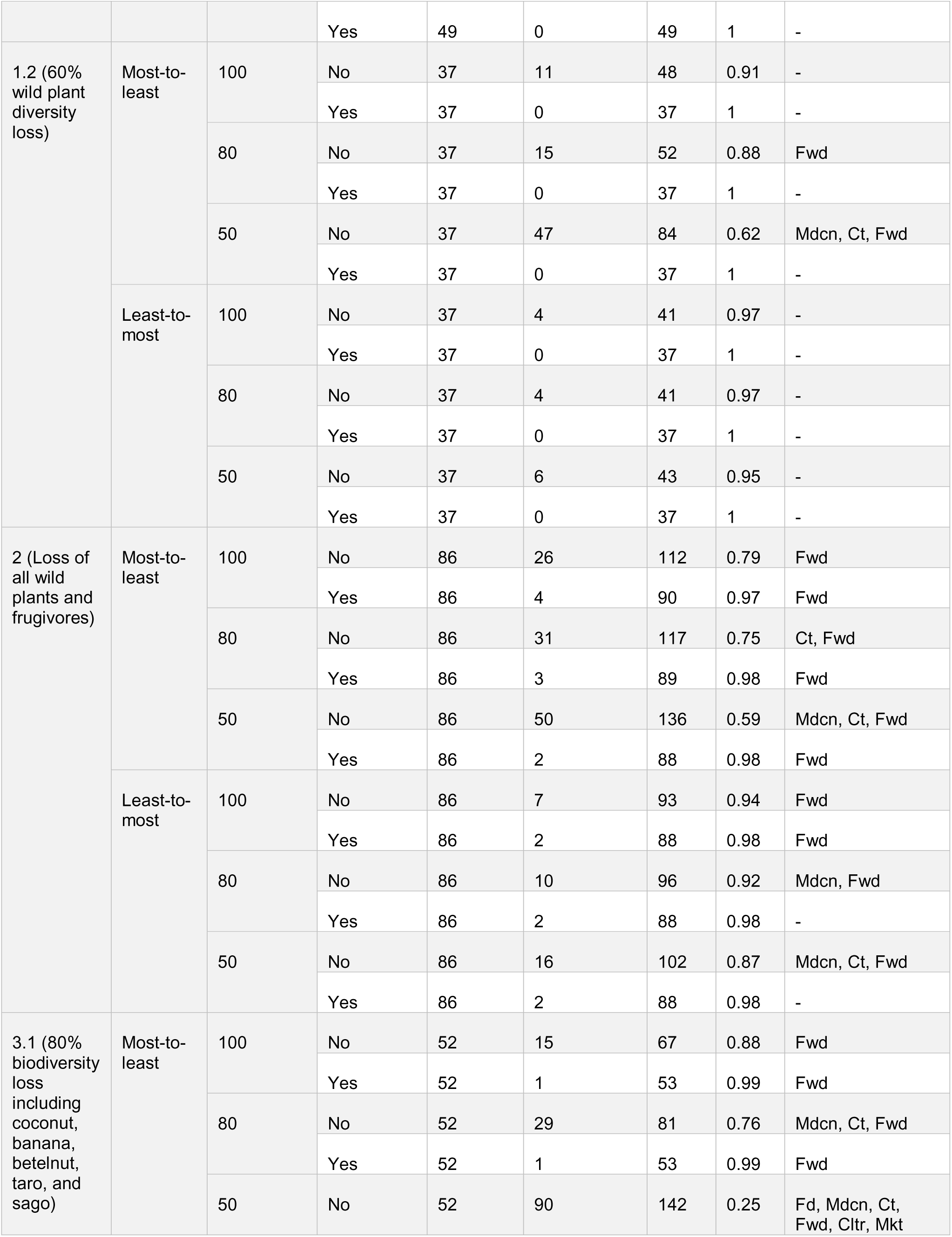
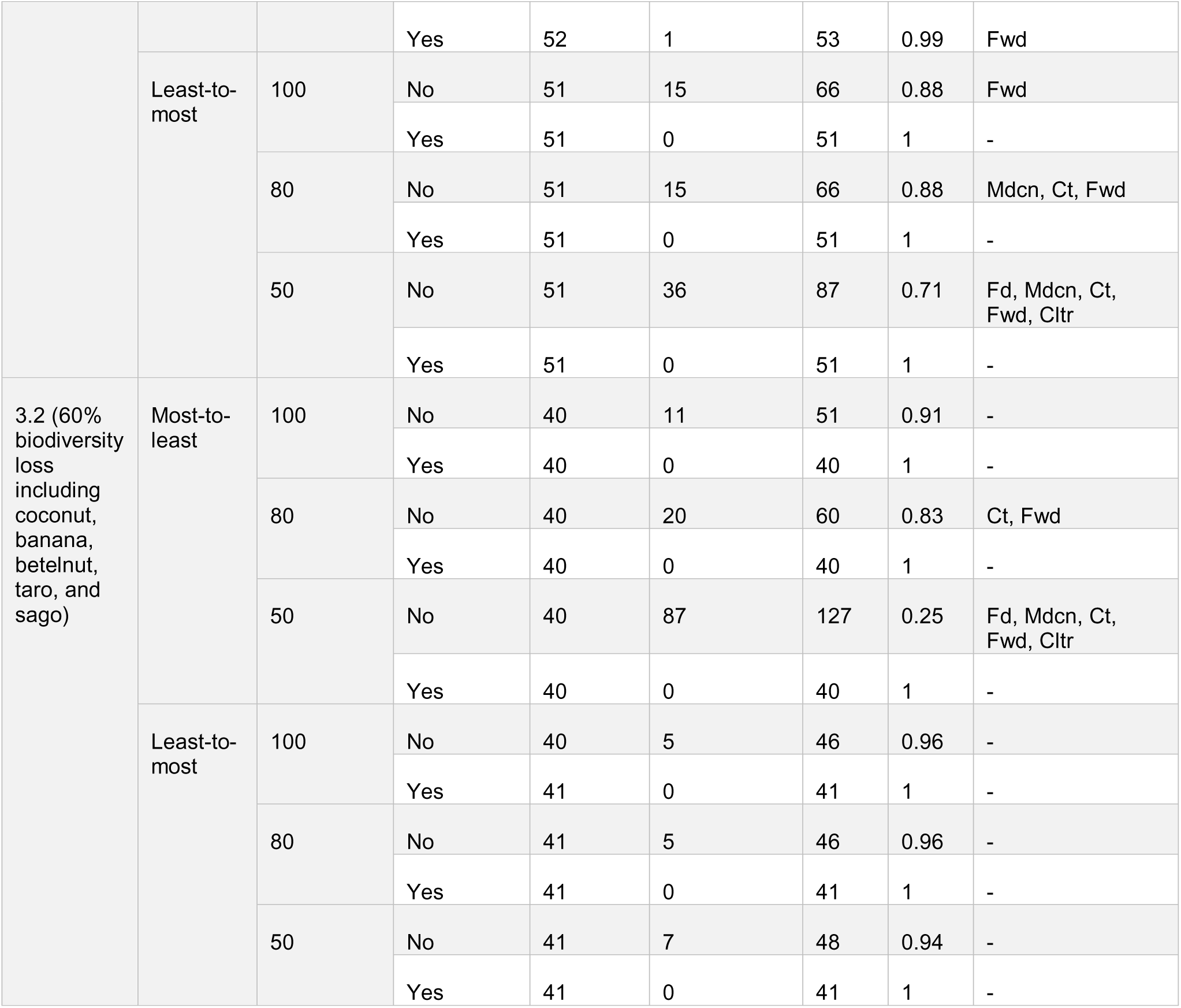
Results of extinction scenarios showing the number of primary, secondary, and total extinctions, network robustness (R100) and the ecosystem services lost. Ecosystem service codes: Fd = Food; Mdcn = Medicine; Ct = Construction and tools; Fwd = Firewood; Cltr = Culture; Mkt = Income from services.

The robustness of our network depended on the order of species extinction and the link loss threshold of nodes within the network. The ‘best-case’ extinction simulations, where species were removed from least-most connected, produced highly robust networks, with collapse occurring only after an average loss of 92% (*R100* = 0.92± 0.02) compared to ‘worst-case’ simulations of 72% of species (*R100* = 0.72 ± 0.06) (Table 2). The worst-case scenarios experienced at least one service loss in 12 of 15 simulations, compared to 6 of 15 in the best-case scenarios.

Link loss thresholds further influenced network robustness. Simulations with a 100% link loss threshold experienced network collapse after approximately 91% of species lost (*R100* = 0.91 ± 0.02), compared to 88% of species under the 80% threshold (*R100* = 0.88 ± 0.03) and 67% below the 50% threshold (*R100* = 0.67 ± 0.08) (Table 2). This indicates that on average across all scenarios a loss of two-thirds of all species may lead to the initial degradation of the network and therefore overall service provision, whereas a loss of 90% of species would be required for a full network collapse and overall ecosystem service loss.

Ecosystem services differed in their sensitivity to species loss. Income from services was the most robust category overall when considering all scenarios and simulations (Fig. 5), requiring essentially all species to be removed before being lost from the network (*R* = 0.999 ±0.001; Table S2). Firewood was the least robust ecosystem service (Fig. 5), with a reduction, functional or complete loss occurring in more than 63% of scenarios (Table 2; Figs. S1-5), and loss from the network after the removal of an average of 60% of species (*R* = 0.60 ± 0.07; Fig. 5; Table 2). Other services, food, medicine, construction and tools, and cultural, showed intermediate vulnerability to species loss, with their loss depending on the environmental change scenario, extinction order, and link loss threshold (Fig. 5; Table 2).

**Figure 5:**
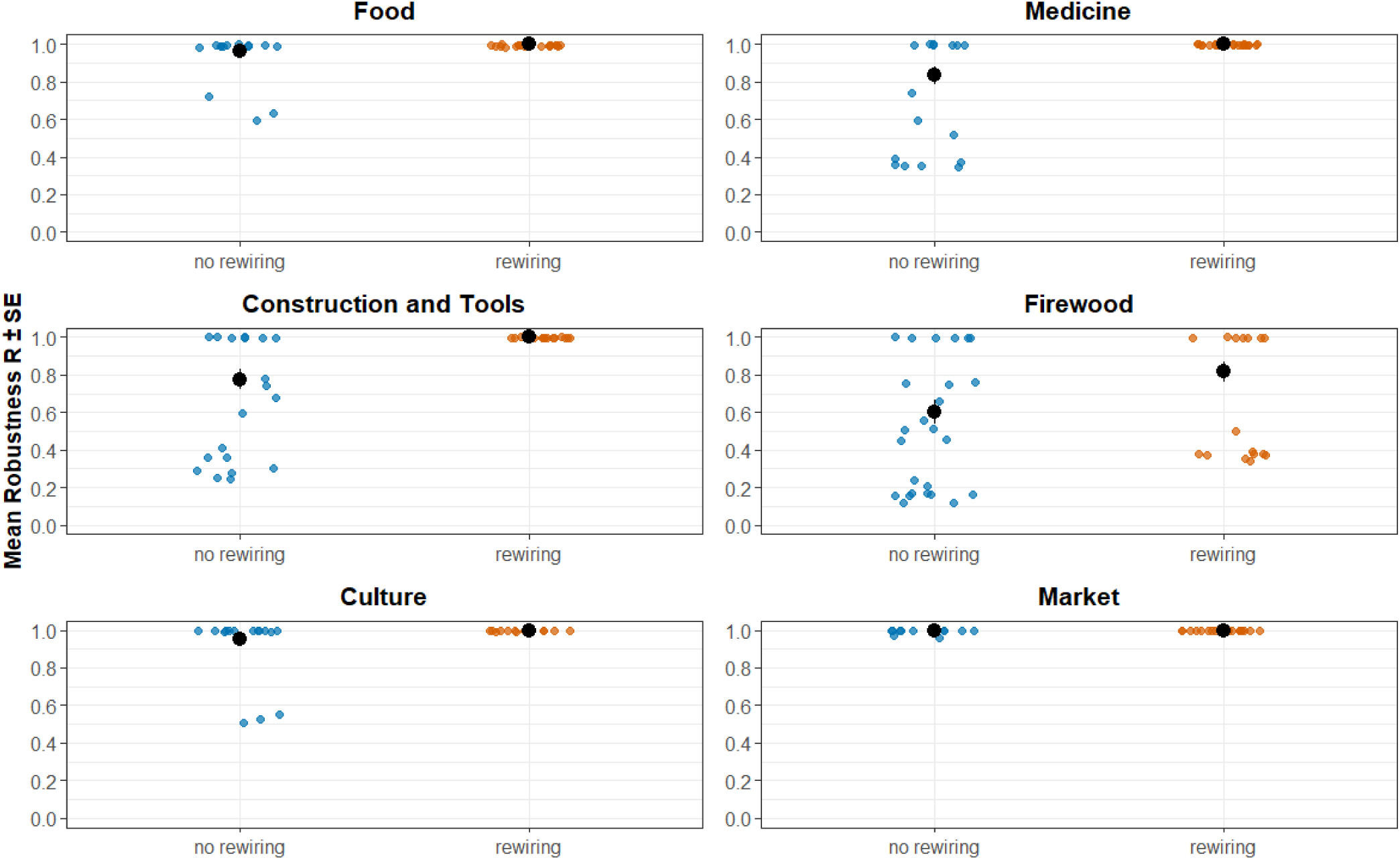
**The average robustness (R) of each ecosystem service category across all extinction simulations**. R indicates the percentage of species within a network that must be removed before an ecosystem service is lost.

### 3.2 The direct and indirect role of species and interaction pathways

Primary extinctions in our socioecological network triggered an average of 22.0 ± 4.1 secondary extinctions per simulation, representing 17.9% of nodes across both network layers (Fig. 3b; Table 2). In three simulations, this was greater than the number of primary extinctions (Table 2), highlighting how species loss can propagate dramatically across socioecological networks to amplify biodiversity loss. The number of secondary extinctions in the network was strongly negatively related to network robustness (regression estimate: -0.008, p < 0.001, R^2^ = 0.99; Fig. 3c), demonstrating that larger extinctions cascades substantially reduce the ability of the network to maintain key ecosystem services.

The magnitude of secondary extinctions triggered by each primary extinction varied widely depending on species identities and network position, ranging from 0 – 63 secondary losses (Fig. S6). The resulting impact this then had on ecosystem service provisioning depended not only on the species directly associated with each service, but also on indirect interaction pathways between biodiversity and ecosystem services. For example, the loss of *Dracontomelon dao* from the plant layer (known locally as Mon) produced the strongest cascading effects across the network (Fig. S6), with extensive interaction pathways connecting its removal to multiple ecosystem services both directly, through its own contributions, and indirectly, by altering interactions among other service-providing plant species through shared interaction partners (Fig. 2). In contrast, other species, such as *Cocos nucifera*, showed considerably fewer secondary extinctions, with minimal impact on the network after their removal. With only one frugivorous interaction, the removal of *C. nucifera* does not propagate through the network (Fig. 2).

Despite these species-specific differences, no single species emerged as disproportionately important for maintaining overall network robustness. No plant or frugivore species recorded a relative importance score greater than 0.4 for any ecosystem service (Table S3), indicating that even the most influential or connected species within the network is important for maintaining robust services in no more than 40% of simulations.

### 3.3 The role of interaction rewiring in ecosystem service robustness and resilience

Running our simulations with interaction rewiring substantially increased socioecological network robustness, allowing the system to withstand the loss of on average 99% of all species before complete collapse (*R100* = 0.99 ± 0.002; Fig. 3a). This was an 18% increase in species lost compared to when rewiring was omitted (Fig. 3a). Consistent with this, rewiring led to a significant reduction in secondary extinctions, with only ∼1 species lost through secondary extinctions (0.7 ± 0.19; Fig. 3b), 21 species less than when rewiring was omitted, resulting in a more robust overall network (Fig. 3b).

Embedded within this increased robustness was a greater resilience of ecosystem service provisioning under environmental change (Fig. 4; 5). The additional robustness provided by interaction rewiring meant that, across all environmental change scenarios, extinction orders, and link-loss thresholds, five out of the six ecosystem services - food, income from services, medicine, construction and tools, and culture - were consistently maintained (Fig. 5; Table 2). The only exception was firewood services, which were still lost in 9 simulations (Fig. 5; Table 2), although this represents a 53% decrease in losses from simulations without rewiring and is largely restricted to the most extreme environmental change simulations (Table 2). Overall, the results indicate that the adaptability of both species and service users following species loss can confer increased resilience.

Interaction rewiring altered the relative importance of species within the socioecological network, including their direct and indirect interactions for maintaining resilient ecosystem services (Table S3). For firewood services, for example, interaction rewiring generally reduced the relative importance scores of plant species (Fig. 6), as the adaptability of species allowed competitors to replace lost interactions. However, several species, *Cocos nucifera* (coconut), *Inocarpus fagifer* (polynesian chestnut), *Metroxylon sagu* (sago), and *Zingiber officinale* (ginger), showed increases in relative importance of up to 0.2, indicating that they contributed to maintaining service provision in 20% more simulations when rewiring was included. Furthermore, after including interaction rewiring within the model, the loss of no frugivorous species had significant effects on ecosystem service robustness, with no frugivores scoring above 0.07 in relative importance for maintaining resilient ecosystem services (Table S3). These patterns suggest that the adaptive capacities of species can shift their functional roles within the socioecological network.

**Figure 6:**
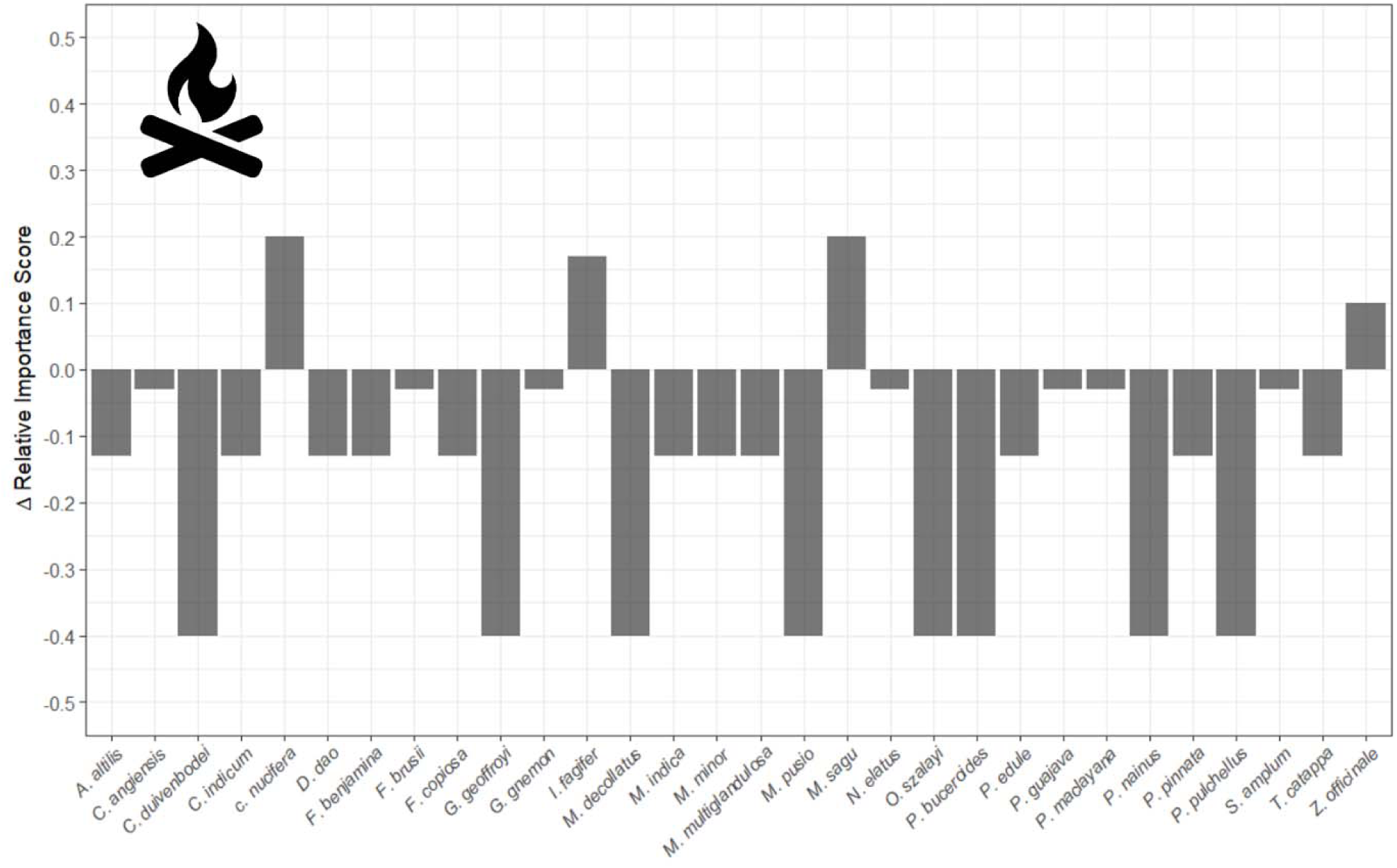
Change in the relative importance scores of important species for maintaining robust firewood services to the local community. The change is directed from no rewiring to including rewiring. Generally, when interaction rewiring was not allowed within the simulations, species scored slightly higher than when interaction rewiring was included (Table S3), however, the relative importance of some species increases when given flexibility in their interaction partners.

## 4. Discussion

Using a socioecological network representing a tropical agricultural system, we evaluated network robustness to simulated species loss with and without interaction rewiring and assessed how primary and secondary species extinctions propagate through interaction pathways and affect ecosystem service provisioning. Our results showed that (1) even without interaction rewiring, ecosystem service provisioning in our network is generally robust and resilient against species loss (Fig. 5; Table 2); (2) ecosystem service provisioning can be threatened through secondary extinctions of species, as interaction pathways can trigger further species loss and functional network collapse, especially under extreme environmental change scenarios (Fig. 2-4); and (3) interaction rewiring enhances ecosystem services resilience against species loss (Fig. 3-5), and alters the relative importance of individual species and their interactions in maintaining ecosystem service provisioning. Together, these results highlight that a wider range of species than just those directly providing ecosystem services should be considered when assessing ecosystem service resilience to maintain human wellbeing and livelihoods. Furthermore, our results suggest that ecosystem services may be more resilient than expected when biodiversity and people are able to adapt to ongoing environmental change.

### 4.1 Ecosystem service robustness and resilience and network adaptability to change

We found that ecosystem services included within our socioecological network, such as food, cultural, and income-providing services, are generally robust against all but the most extreme species loss projections under future environmental change; and that if allowed to adapt, socioecological functioning can be robustly maintained. These results align with previous ecological studies showing that interaction rewiring enhances the robustness and stability of mutualistic plant-pollinator networks (Kaiser-Bunbury et al., 2010; Vizentin-Bugoni et al., 2020), yet rewiring has rarely been considered in socioecological networks or ecosystem service contexts (Keyes et al., 2021). Our work suggests that if ecological and local communities can shift their dependence or use alternative natural resources, then ecosystem service provisioning may be more resilient to projected species loss under future environmental change than expected. However, it is important to note that although overall service provision may be retained, loss of food sources, medicinal plants, and crops and wild plants to sell at markets can still reduce the diversity of nutrients, treatments available, and income available to local communities (Alves & Rosa, 2007; Asigbaase et al., 2023; Linhares et al., 2023; Nguyen & Qaim, 2025). Therefore, whilst our results indicate that adaptability can help maintain overall service delivery, preventative measures will still be required to protect vulnerable species and ecosystem functions that contribute to human wellbeing and livelihoods (IPBES 2019).

Rewiring increased the number of species our network could lose before functionally collapsing and losing overall ecosystem service provisioning, with adaptive interactions allowing network structures to persist after the loss of nearly 20% more species compared with scenarios without rewiring (Fig. 3a). These results demonstrate how incorporating interaction rewiring into ecosystem service assessments could help evaluate the adaptive potential of socioecological systems, highlighting when adaption may be an effective protective strategy and where it is unlikely to provide substantial benefits. Adaptability may, for example, be insufficient in communities that rely on specific plants to supply key nutrients. For example, in a smallholder farming community in Nepal, carrot flowers are pollinated almost exclusively by flies and provide an important source of vitamin A to the local community. This suggests that, in this context, ecological and social adaptability may not be sufficient to buffer against species loss, which reduces pollination services underpinning key nutrients (Timberlake et al., 2022). The adaptive potential of our network could also reflect the strong local knowledge of plants and their uses in the Ohu community, which enhances the resilience of subsistence ecosystem services by enabling local communities to adjust to ecological change (Cámara-Leret et al., 2019).

However, in Papau New Guinea and worldwide, there are growing concerns about the loss of local and indigenous knowledge and migration from local communities weakening food and medicinal systems, potentially reducing the adaptive capacity of future generations amid increasing reliance on natural, physical and social capitals (Cámara-Leret, Raes, et al., 2019; Cámara-Leret & Bascompte, 2021; Hazenbosch et al., 2022; Kik et al., 2021; Powell et al., 2017; Zimmermann et al., 2024). It is also important to note that whilst adaptability may provide short-term resilience in ecosystem service provisioning, pressure may be increased on alternative species over time, reducing general ecosystem services long-term (Tilman et al., 1994). Therefore, ecological and socioecological adaptability to change, and its consequences, is complex and requires careful consideration when assessing short vs long-term ecosystem service resilience and informing management strategies. The inclusion of ecosystem services as network nodes enabled us to quantify the vulnerability of individual services to collapse under species loss, and whose maintained provisioning may need active management efforts to protect (Fig. 1a; 2). For example, the results of our extinction modelling indicate that firewood ecosystem services are the most vulnerable in the Ohu community. This is to be expected, given that the local community generally relies on a single dominant species (*Pometia pinnata*) for firewood provisions, with less flexibility to use other species compared to other services (see Stanworth et al., 2026). By identifying firewood services as vulnerable, this can facilitate targeted modelling of firewood flows in local PNG communities to better understand how firewood is used and the regeneration of firewood harvests. This would allow for more nuanced predictions about how firewood supplies may change under species losses resulting in better informed land management and decision making (Nuberg 2015).

### 4.2 The role of species and interaction pathways

Our results highlight the role of indirect interaction pathways in supporting robust and resilient ecosystem service provisioning. Across extinction scenarios, the loss of *Dracontomelon dao* consistently resulted in the highest number of secondary extinctions. *D. dao* provides all six ecosystem services measured in this study. Thus, whilst its loss demonstrates how the removal of a key species can propagate through a network to affect ecosystem service flows (Fig. 2), its central role in supporting service provisioning is largely predictable. However, the impact of losing other species on ecosystem service provisioning was less predictable. For example, the loss of *Metroxylon sagu* (sago) and *Cocos nucifera* had significant impacts on the resilience of firewood services, despite neither species directly providing firewood to the local community. The initial loss of each species and the subsequent adaptation by both biodiversity and local farmers increased the importance of indirect interaction pathways that connect *M. sagu and C. nucifera* to other species that provide firewood through shared frugivorous interaction partners. These findings demonstrate that species considered important for directly providing ecosystem services may differ from those that underpin service robustness through their indirect interaction pathways. Therefore, incorporating interaction rewiring further reveals novel species interaction pathways that are important for maintaining resilient ecosystem services. Consequently, failing to incorporate ecological and socioecological adaptations into species loss modelling can lead to both over- and underestimating the roles of species and their indirect interactions in maintaining ecosystem services (Kaiser-Bunbury et al., 2010; Keyes et al., 2021).

Furthermore, although the loss of frugivores had cascading effects through the socioecological network when interaction rewiring was not incorporated, the loss of no frugivorous species significantly affected ecosystem service resilience through their indirect interaction pathways after interactions were shuffled in response to species loss. Frugivores are known to adapt to declining resource availability (Costa et al., 2028), and our study demonstrates how this adaptive capacity translates into resilient ecosystem services through maintaining ecological functioning and indirect pathways to service delivery. These results highlight the extent to which the reorganisation of ecological and socioecological interactions following primary species loss can play a role in sustaining ecosystem service flows. For example, rewiring can strengthen existing ecological and socioecological interactions or generate new ones, as frugivores and farmers are forced to replace or shift reliance on natural resources after species loss.

Consequently, peripheral species that do not directly provide any ecosystem services, or that initially occupy weakly connected positions in ecological networks, may become more central to ecosystem service flows as realised links change and new indirect connections to ecosystem services are created, or vice versa (Costa et al., 2018; Goldstein & Zych, 2016; Timóteo et al., 2016). Therefore, focusing solely on species that directly provide ecosystem services can lead to underestimation of the importance of other species and their interaction pathways in indirectly supporting services, or, equally, the overestimation of the role of some species at the expense of highlighting others (Keyes et al., 2021; Rey et al., 2024). This insight is key for land management strategies that seek to support dual biodiversity and ecosystem service outcomes, as sustaining human well-being and livelihoods may require a larger pool of species than is suggested by traditional ecosystem service assessments that do not account for flexibility in species’ interactions in service resilience (Kaiser-Bunbury & Blüthgen, 2015; Keyes et al., 2021).

### 4.3 Limitations and next steps

Our work acts as a proof-of-concept demonstrating that adaptability can be incorporated into ecosystem service assessments by using interaction rewiring in socioecological network models and highlighting the importance of indirect interaction pathways in determining ecosystem service resilience. It is therefore worth noting that, whilst we have tried to account for different magnitudes of environmental change within our simulations, our extinction scenarios and interaction rewiring are still simplified and do not account for iterative changes such as introduced species or the effect of colonisation (Bennett & Reyers, 2024; Keyes et al., 2021; Marjakangas et al., 2025). In addition, the ecological interaction rewiring probabilities in our extinction simulations was calculated using a latent-trait probability model that infers how interactions will reshuffle based on existing network structure (Terry & Lewis, 2020), and does not directly incorporate measures of species morphological or phenological specialisation that could improve estimates of species adaptability (Ducatez et al., 2020). Furthermore, by using interviews to compile species interaction data, it is likely that we are missing rare or nocturnal interactions (Carlo & Morales, 2016; Martinez & Pires, 2024; Ong et al., 2021), and that some reported frugivorous interactions reflect disservices, such as frugivores that consume cultivated plants or destroy seeds that would otherwise be consumed or sold by farmers (Dracxler & Kissling, 2022; Stanworth et al., 2026). Considering these factors, our results should be interpreted as guidance on how species losses can influence the robustness of ecosystem services, and how both species-level and human adaptability can be incorporated into assessments of ecosystem service resilience – rather than to directly influence management decisions. To improve the accuracy of such assessments and generate outputs that can more reliably inform management decisions, practitioners may wish to integrate behavioural or trait-based data to better capture species interaction flexibility (Bachmann & Drossel, 2025; Dehling et al., 2016). They may also need to consider the type of interactions (e.g. mutualistic vs antagonistic) as antagonistic interactions could affect the robustness and resilience of ecosystem services through their presence, loss or emergence (Briggs et al., 2019).

When incorporating interview data, practitioners could adopt a more iterative approach to better capture dynamic human-nature relationships shaped by preferences (Bennett & Reyers, 2024), and elicit participant’s adaptive response. One way to achieve this could be to undertake two-stage interviews where initial interviews provide information for socioecological interactions connecting biodiversity to ecosystem services, and subsequent interviews explore how participants would respond to projected species loss and primary and secondary extinctions, such as by substituting alternative natural or non-natural resources. These context-specific socioecological data would capture flexibility and nuances in human preferences that shape ecosystem service use and offer a more detailed understanding of service resilience grounded by both ecological and socioecological adaptation to change (Cámara-Leret et al., 2019).

## 5. Concluding remarks

Our study quantified the resilience of ecosystem service provisioning under a range of environmental change scenarios, showing that when biodiversity and local communities can adapt, ecosystem services can remain resilient to species losses. We found that vulnerability is not always driven solely by the loss of direct service providers; instead, biodiversity often plays an important indirect role in maintaining robust and resilient ecosystem services.

Although our results are based on a smallholder farming community in Papau New Guinea, our framework is broadly applicable for assessing how species losses affect ecosystem services and the indirect interaction pathways that underpin resilient subsistence systems. For example, reef-dependent coastal and pastoral rangeland communities have both been identified as vulnerable to species loss under future environmental changes (Hughes et al., 2017; Reid et al., 2014), and our approach could offer deeper insight into how these losses cascade through ecosystem service networks. By improving our understanding of the indirect contributions that species play in supporting human wellbeing and livelihoods, we can better anticipate the outcome of future climatic or socioecological change for the provision of local ecosystem services. This can enable land management strategies to move beyond prioritising primary service providers, to considering all species that support the resilience of these services and whose loss could propagate through interaction networks and ultimately diminish or eliminate key ecosystem services.

## Author contributions

Anna Stanworth, Kelvin S.-H. Peh and Rebecca J. Morris conceived the ideas. Kiole Imale facilitated and led data collection in Ohu, remotely supported by Anna Stanworth; Anna Stanworth did data analysis, led writing of the manuscript and made figures. Anna Stanworth, Harry E.R. Shepherd, Kelvin S.-H. Peh and Rebecca J. Morris all contributed to ongoing discussion and drafts of the manuscript. All authors gave final approval for publication.

## Statement on Inclusion

Our study brings together authors based in the UK and Papua New Guinea, where the data for this study was collected, and authors from Papua New Guinea led data collection. Efforts were made to consider previous work published by the local research centre in Papua New Guinea, including studies carried out in the specific area in which the study is based.

## Acknowledgments

We are in debt to the community of Ohu who have shared information about their livelihoods, and local ecological knowledge with us. We are very thankful for the support from staff at the New Guinea Binatang Research Centre, specifically Kiole Imale, Brus Isua and Anne M. Anji who facilitated and led data collection for the data used in this study. We are also thankful to note takers that enabled data collection, Agatha Teisua for early project support, and ongoing project support from Dr Francesca Dem, Professor Vojtěch Novotný, and Dr Pagi Toko. This work was supported by the Natural Environment Research Council (NERC) (grant number NE/2007210/1) through the INSPIRE DTP, and the RGS-IBG Postgraduate Research Award, Royal Geographical Society (with IBG).

## Data Availability Statement

All data are publicly available at https://doi.org/10.5281/zenodo.17515454.

