## Supplementary Materials for "Socioecological network adaptability enhances ecosystem service resilience in a tropical agricultural system"

Appendix S1: Edge list of interactions included in the tripartite socioecological network composed of two layers of interactions; ecological frugivorous interactions and socio-ecological plant-benefit interactions – *Stanworth_et_al_frugivory_plant_ES_data.csv* doi: https://doi.org/10.5281/zenodo.17515454

Table S1: Summary of extinction scenario simulations.

| Simulation number | Simulation description | Simulation code |
| --- | --- | --- |
| 1 | 1.1 80% loss of wild plant diversity under RCP 8.5; Most to least connected; 100% link loss | 1.1ml100 / 1.1ml100_rewired |
| 2 | 1.1 80% loss of wild plant diversity under RCP 8.5; Most to least connected; 80% link loss | 1.1ml80 / 1.1ml80_rewired |
| 3 | 1.1 80% loss of wild plant diversity under RCP 8.5; Most to least connected; 80% link loss | 1.1ml50 / 1.1ml50_rewired |
| 4 | 1.1 80% loss of wild plant diversity under RCP 8.5; Least to most connected; 100% link loss | 1.1lm100 / 1.1lm100_rewired |
| 5 | 1.1 80% loss of wild plant diversity under RCP 8.5; Least to most connected; 80% link loss | 1.1lm80 / 1.1lm80_rewired |
| 6 | 1.1 80% loss of wild plant diversity under RCP 8.5; Least to most connected; 50% link loss | 1.1lm50 / 1.1lm50_rewired |
| 7 | 1.2 60% loss of wild plant diversity under RCP 2.6; Most to least connected; 100% link loss | 1.2ml100 / 1.2ml100_rewired |
| 8 | 1.2 60% loss of wild plant diversity under RCP 2.6; Most to least connected; 80% link loss | 1.2ml80 / 1.2ml80_rewired |
| 9 | 1.2 60% loss of wild plant diversity under RCP 2.6; Most to least connected; 50% link loss | 1.2ml50 / 1.2ml50_rewired |
| 10 | 1.2 60% loss of wild plant diversity under RCP 2.6; Least to most connected; 100% link loss | 1.2lm100 / 1.2lm100_rewired |
| 11 | 1.2 60% loss of wild plant diversity under RCP 2.6; Least to most connected; 80% link loss | 1.2lm80 / 1.2lm80_rewired |
| 12 | 1.2 60% loss of wild plant diversity under RCP 2.6; Least to most connected; 50% link loss | 1.2lm50 / 1.2lm50_rewired |
| 13 | 2 Loss of biodiversity under forest loss for garden expansion; Most to least connected; 100% link loss | 2ml100 / 2ml100_rewired |
| 14 | 2 Loss of biodiversity under forest loss for garden expansion; Most to least connected; 80% link loss | 2ml80 / 2ml80_rewired |
| 15 | 2 Loss of biodiversity under forest loss for garden expansion; Most to least connected; 50% link loss | 2ml50 / 2ml50_rewired |
| 16 | 2 Loss of biodiversity under forest loss for garden expansion; Least to most connected; 100% link loss | 2lm100 / 2lm100_rewired |
| 17 | 2 Loss of biodiversity under forest loss for garden expansion; Least to most connected; 80% link loss | 2lm80 / 2lm80_rewired |
| 18 | 2 Loss of biodiversity under forest loss for garden expansion; Least to most connected; 50% link loss | 2lm50 / 2lm50_rewired |
| 19 | 3.1 Tandem loss of 80% of wild plant species richness under RCP 8.5, loss of species to disease (coconut, banana, betelnut, taro), and loss of species to overharvesting (sago); Most to least connected; 100% link loss | 3.1ml100 / 3.1ml100_rewired |
| 20 | 3.1 Tandem loss of 80% of wild plant species richness under RCP 8.5, loss of species to disease (coconut, banana, betelnut, taro), and loss of species to overharvesting (sago); Most to least connected; 80% link loss | 3.1ml80 / 3.1ml80_rewired |
| 21 | 3.1 Tandem loss of 80% of wild plant species richness under RCP 8.5, loss of species to disease (coconut, banana, betelnut, taro), and loss of species to overharvesting (sago); Most to least connected; 50% link loss | 3.1ml50 / 3.1ml50_rewired |
| 22 | 3.1 Tandem loss of 80% of wild plant species richness under RCP 8.5, loss of species to disease (coconut, banana, betelnut, taro), and loss of species to overharvesting (sago); Least to most connected; 100% link loss | 3.1lm100 / 3.1lm100_rewired |
| 23 | 3.1 Tandem loss of 80% of wild plant species richness under RCP 8.5, loss of species to disease (coconut, banana, betelnut, taro), and loss of species to overharvesting (sago); Least to most connected; 80% link loss | 3.1lm80 / 3.1lm80_rewired |
| 24 | 3.1 Tandem loss of 80% of wild plant species richness under RCP 8.5, loss of species to disease (coconut, banana, betelnut, taro), and loss of species to overharvesting (sago); Least to most connected; 50% link loss | 3.1lm50 / 3.1lm50_rewired |
| 25 | 3.2 Tandem loss of 60% of wild plant species richness under RCP 2.6, loss of species to disease (coconut, banana, betelnut, taro), and loss of species to overharvesting (sago); Most to least connected; 100% link loss | 3.2ml100 / 3.2ml100_rewired |
| 26 | 3.2 Tandem loss of 60% of wild plant species richness under RCP 2.6, loss of species to disease (coconut, banana, betelnut, taro), and loss of species to overharvesting (sago); Most to least connected; 80% link loss | 3.2ml80 / 3.2ml80_rewired |
| 27 | 3.2 Tandem loss of 60% of wild plant species richness under RCP 2.6, loss of species to disease (coconut, banana, betelnut, taro), and loss of species to overharvesting (sago); Most to least connected; 50% link loss | 3.2ml50 / 3.2ml50_rewired |
| 28 | 3.2 Tandem loss of 60% of wild plant species richness under RCP 2.6, loss of species to disease (coconut, banana, betelnut, taro), and loss of species to overharvesting (sago); Least to most connected; 100% link loss | 3.2lm100 / 3.2lm100_rewired |
| 29 | 3.2 Tandem loss of 60% of wild plant species richness under RCP 2.6, loss of species to disease (coconut, banana, betelnut, taro), and loss of species to overharvesting (sago); Least to most connected; 80% link loss | 3.2lm80 / 3.2lm80_rewired |
| 30 | 3.2 Tandem loss of 60% of wild plant species richness under RCP 2.6, loss of species to disease (coconut, banana, betelnut, taro), and loss of species to overharvesting (sago); Least to most connected; 50% link loss | 3.2lm50 / 3.2lm50_rewired |

Table S2: Individual ecosystem service robustness results. The following shorthand refers to each ecosystem service category: Fd – Food, Mdcn – Medicine, Ct – Construction and Tools, Fwd – Firewood, Cltr – Culture, Mkt – income from services. R = robustness.

| Simulation code | R100 | Fd R | | Mdcn R | | Ct R | Fwd R | Cltr R | Mkt R |
| --- | --- | --- | --- | --- | --- | --- | --- | --- | --- |
| 1.1ml100 | 0.88 | | 1 | | 1 | 1 | 0.45 | 1 | 1 |
| 1.1ml100_rewired | 0.99 | | 1 | | 1 | 1 | 0.37 | 1 | 1 |
| 1.1ml80 | 0.85 | | 1 | | 1 | 1 | 0.16 | 1 | 1 |
| 1.1ml80_rewired | 0.99 | | 1 | | 1 | 1 | 0.37 | 1 | 1 |
| 1.1ml50 | 0.6 | | 1 | | 0.37 | 0.25 | 0.12 | 1 | 1 |
| 1.1ml50_rewired | 0.99 | | 1 | | 1 | 1 | 0.35 | 1 | 1 |
| 1.1lm100 | 0.96 | | 1 | | 1 | 1 | 1 | 1 | 1 |
| 1.1lm100_rewired | 1 | | 1 | | 1 | 1 | 1 | 1 | 1 |
| 1.1lm80 | 0.96 | | 1 | | 1 | 1 | 1 | 1 | 1 |
| 1.1lm80_rewired | 1 | | 1 | | 1 | 1 | 1 | 1 | 1 |
| 1.1lm50 | 0.94 | | 1 | | 1 | 1 | 1 | 1 | 1 |
| 1.1lm50_rewired | 1 | | 1 | | 1 | 1 | 1 | 1 | 1 |
| 1.2ml100 | 0.91 | | 1 | | 1 | 1 | 1 | 1 | 1 |
| 1.2ml100_rewired | 1 | | 1 | | 1 | 1 | 1 | 1 | 1 |
| 1.2ml80 | 0.88 | | 1 | | 1 | 1 | 0.16 | 1 | 1 |
| 1.2ml80_rewired | 1 | | 1 | | 1 | 1 | 1 | 1 | 1 |
| 1.2ml50 | 0.62 | | 1 | | 0.35 | 0.25 | 0.15 | 1 | 1 |
| 1.2ml50_rewired | 1 | | 1 | | 1 | 1 | 1 | 1 | 1 |
| 1.2lm100 | 0.97 | | 1 | | 1 | 1 | 1 | 1 | 1 |
| 1.2lm100_rewired | 1 | | 1 | | 1 | 1 | 1 | 1 | 1 |
| 1.2lm80 | 0.97 | | 1 | | 1 | 1 | 1 | 1 | 1 |
| 1.2lm80_rewired | 1 | | 1 | | 1 | 1 | 1 | 1 | 1 |
| 1.2lm50 | 0.95 | | 1 | | 1 | 1 | 1 | 1 | 1 |
| 1.2lm50_rewired | 1 | | 1 | | 1 | 1 | 1 | 1 | 1 |
| 2ml100 | 0.79 | | 1 | | 1 | 1 | 0.66 | 1 | 1 |
| 2ml100_rewired | 0.97 | | 1 | | 1 | 1 | 0.5 | 1 | 1 |
| 2ml80 | 0.75 | | 1 | | 1 | 0.68 | 0.24 | 1 | 1 |
| 2ml80_rewired | 0.98 | | 1 | | 1 | 1 | 0.39 | 1 | 1 |
| 2ml50 | 0.59 | | 1 | | 0.36 | 0.29 | 0.21 | 1 | 1 |
| 2ml50_rewired | 0.98 | | 1 | | 1 | 1 | 0.34 | 1 | 1 |
| 2lm100 | 0.94 | | 1 | | 1 | 1 | 0.76 | 1 | 1 |
| 2lm100_rewired | 0.98 | | 1 | | 1 | 1 | 1 | 1 | 1 |
| 2lm80 | 0.92 | | 1 | | 1 | 0.78 | 0.75 | 1 | 1 |
| 2lm80_rewired | 0.98 | | 1 | | 1 | 1 | 1 | 1 | 1 |
| 2lm50 | 0.87 | | 1 | | 0.74 | 0.74 | 0.76 | 1 | 1 |
| 2lm50_rewired | 0.98 | | 1 | | 1 | 1 | 1 | 1 | 1 |
| 3.1ml100 | 0.88 | | 1 | | 1 | 1 | 0.45 | 1 | 1 |
| 3.1ml100_rewired | 0.99 | | 1 | | 1 | 1 | 0.38 | 1 | 1 |
| 3.1ml80 | 0.76 | | 1 | | 0.52 | 0.36 | 0.17 | 1 | 1 |
| 3.1ml80_rewired | 0.99 | | 1 | | 1 | 1 | 0.38 | 1 | 1 |
| 3.1ml50 | 0.25 | | 0.6 | | 0.35 | 0.3 | 0.15 | 0.5 | 0.96 |
| 3.1ml50_rewired | 0.99 | | 1 | | 1 | 1 | 0.38 | 1 | 1 |
| 3.1lm100 | 0.88 | | 1 | | 1 | 1 | 0.56 | 1 | 1 |
| 3.1lm100_rewired | 1 | | 1 | | 1 | 1 | 1 | 1 | 1 |
| 3.1lm80 | 0.88 | | 1 | | 0.59 | 0.59 | 0.51 | 1 | 1 |
| 3.1lm80_rewired | 1 | | 1 | | 1 | 1 | 1 | 1 | 1 |
| 3.1lm50 | 0.71 | | 0.71 | | 0.39 | 0.41 | 0.51 | 0.56 | 1 |
| 3.1lm50_rewired | 1 | | 1 | | 1 | 1 | 1 | 1 | 1 |
| 3.2ml100 | 0.91 | | 1 | | 1 | 1 | 1 | 1 | 1 |
| 3.2ml100_rewired | 1 | | 1 | | 1 | 1 | 1 | 1 | 1 |
| 3.2ml80 | 0.83 | | 1 | | 1 | 0.36 | 0.17 | 1 | 1 |
| 3.2ml80_rewired | 1 | | 1 | | 1 | 1 | 1 | 1 | 1 |
| 3.2ml50 | 0.25 | | 0.64 | | 0.35 | 0.28 | 0.12 | 0.53 | 1 |
| 3.2ml50_rewired | 1 | | 1 | | 1 | 1 | 1 | 1 | 1 |
| 3.2lm100 | 0.96 | | 1 | | 1 | 1 | 1 | 1 | 1 |
| 3.2lm100_rewired | 1 | | 1 | | 1 | 1 | 1 | 1 | 1 |
| 3.2lm80 | 0.96 | | 1 | | 1 | 1 | 1 | 1 | 1 |
| 3.2lm80_rewired | 1 | | 1 | | 1 | 1 | 1 | 1 | 1 |
| 3.2lm50 | 0.94 | | 1 | | 1 | 1 | 1 | 1 | 1 |
| 3.2lm50_rewired | 1 | | 1 | | 1 | 1 | 1 | 1 | 1 |

.
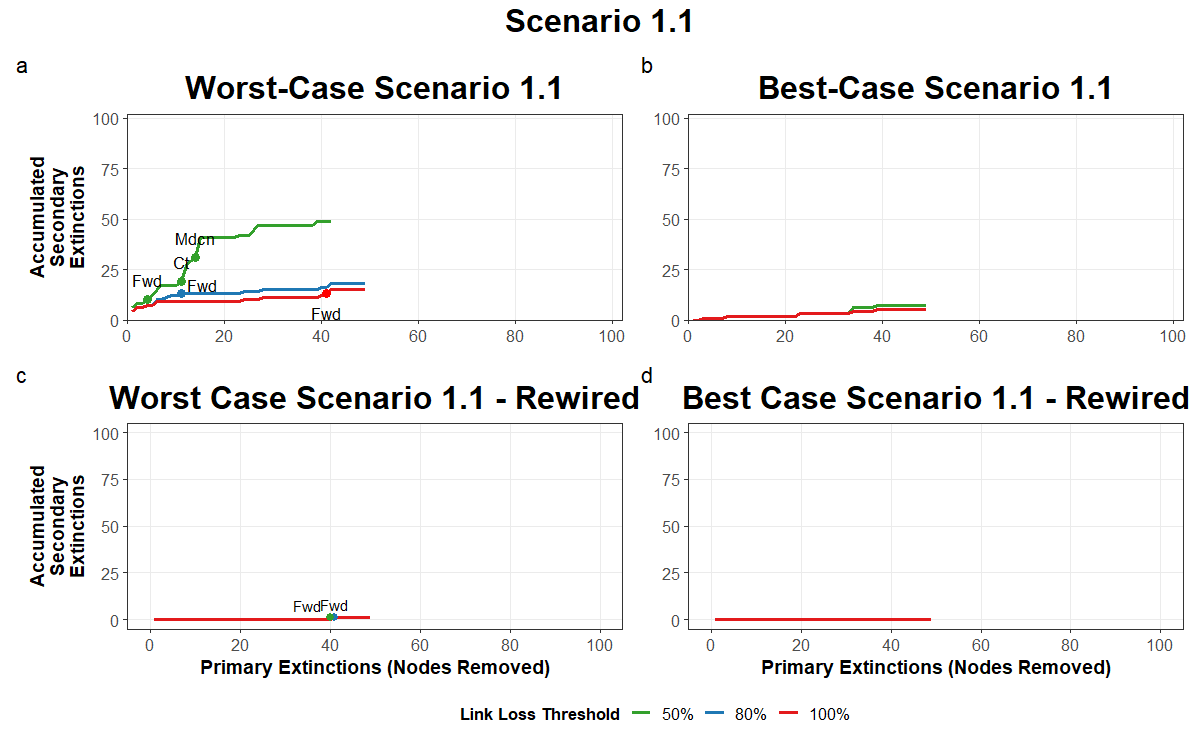

Figure S1: Accumulated secondary extinction graphs for Scenario 1.1. Graphs show secondary extinctions after primary extinction simulations with both interaction rewiring and no interaction rewiring. 50%, 80% and 100% link loss thresholds represent a reduction, functional loss, and complete loss of ecosystem service provisioning for the local community. Points at which an ecosystem service is reduced, functionally lost or completely lost are labeled with fd (food), mkt (income from services), mdcn (medicine), ct (construction and tools), cltr (culture) and fwd (firewood). Under the allocation of interaction rewiring secondary extinctions follow the same or similar pattern under all link loss thresholds, hence the lines have been layered and only the 80% link loss threshold line is visible in rewired graphs.

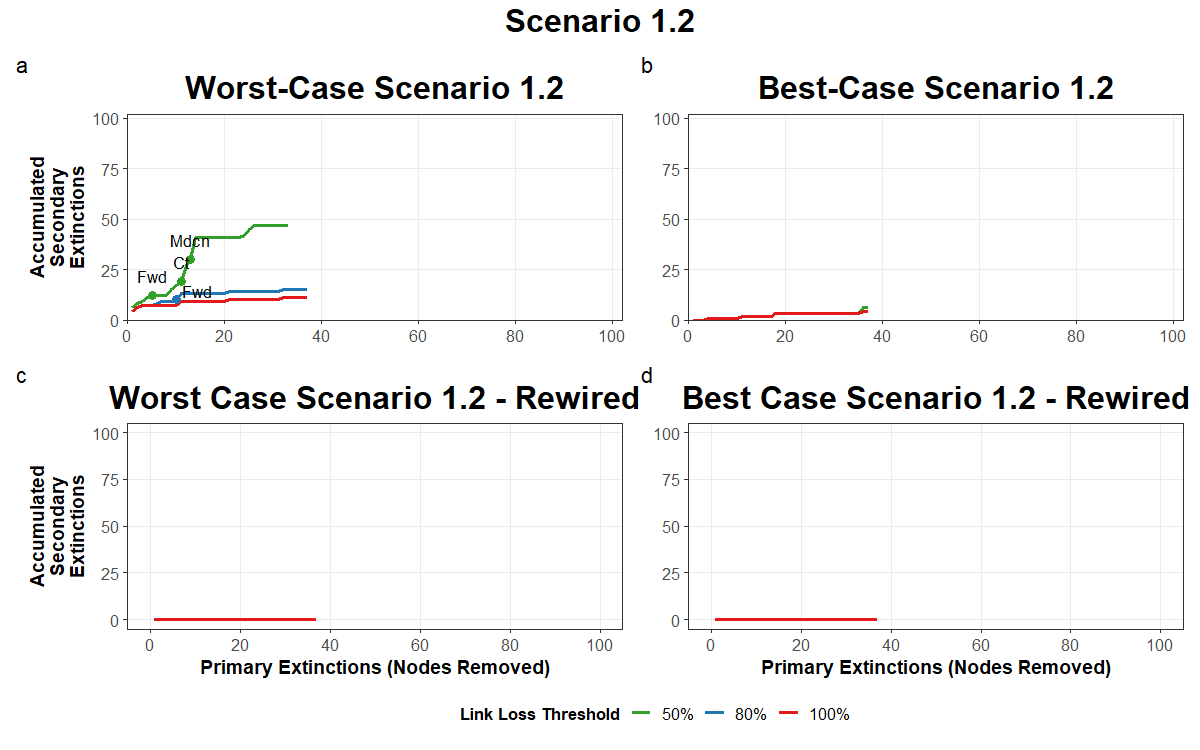

Figure S2: Accumulated secondary extinction graphs for Scenario 1.2. See caption of Fig. S1 for full description of graphs.

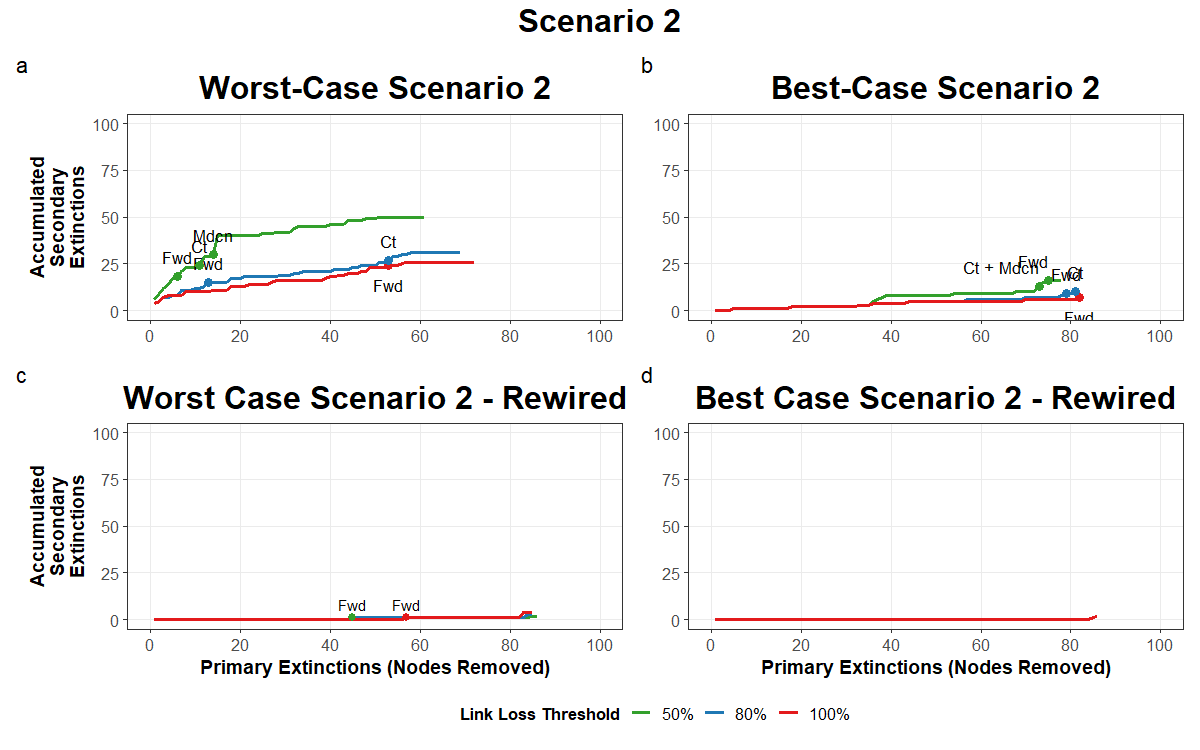

Figure S3: Accumulated secondary extinction graphs for Scenario 2. See caption of Fig. S1 for full description of graphs.

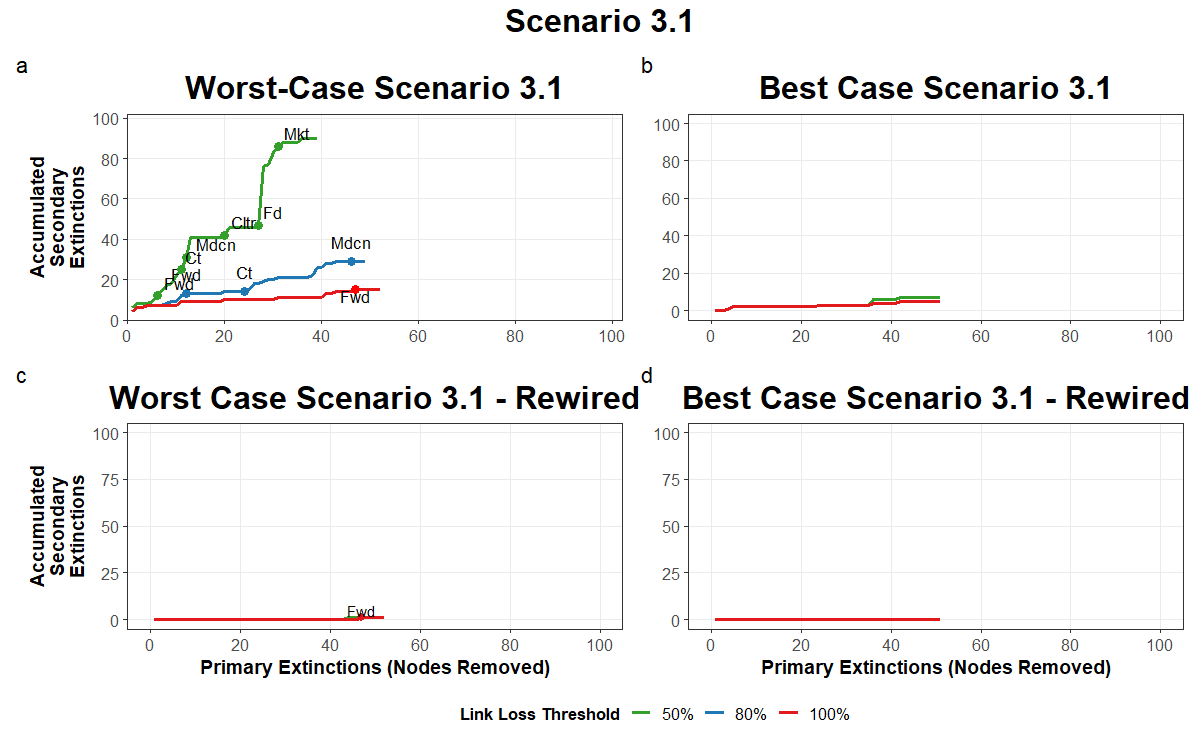

Figure S4: Accumulated secondary extinction graphs for Scenario 3.1. See caption of Fig. S1 for full description of graphs.

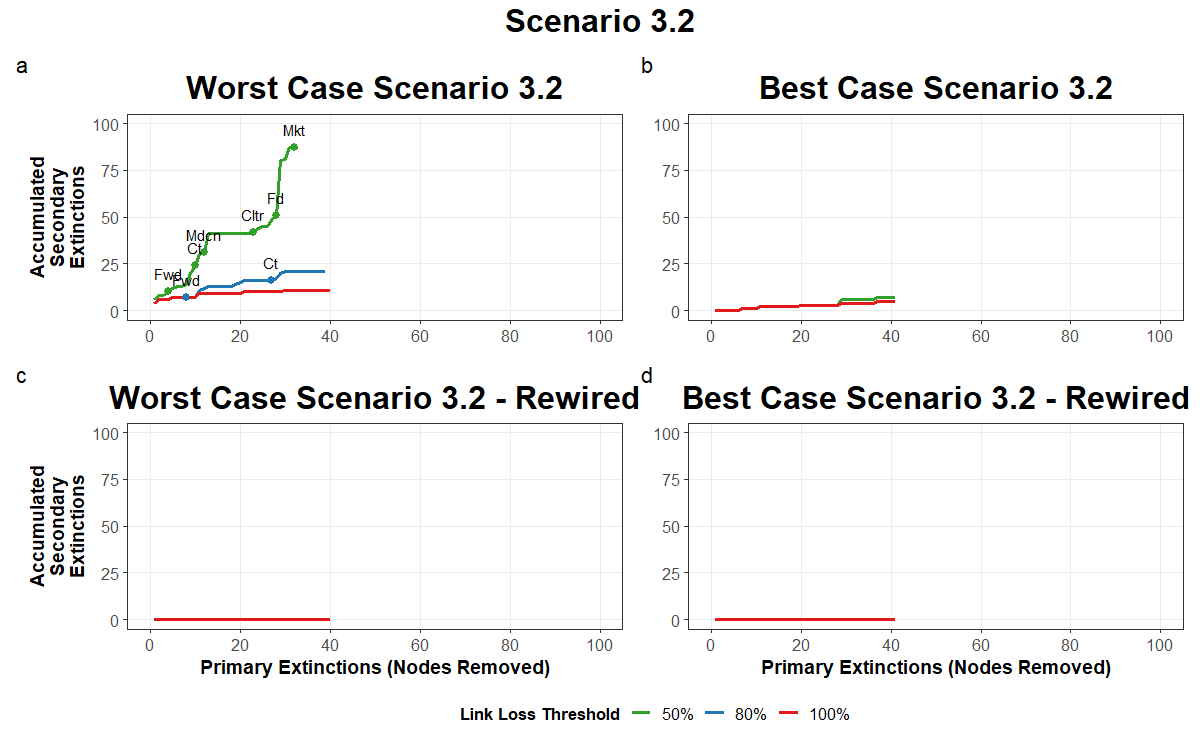

Figure S5: Accumulated secondary extinction graphs for Scenario 3.2. See caption of Fig. S1 for full description of graphs.

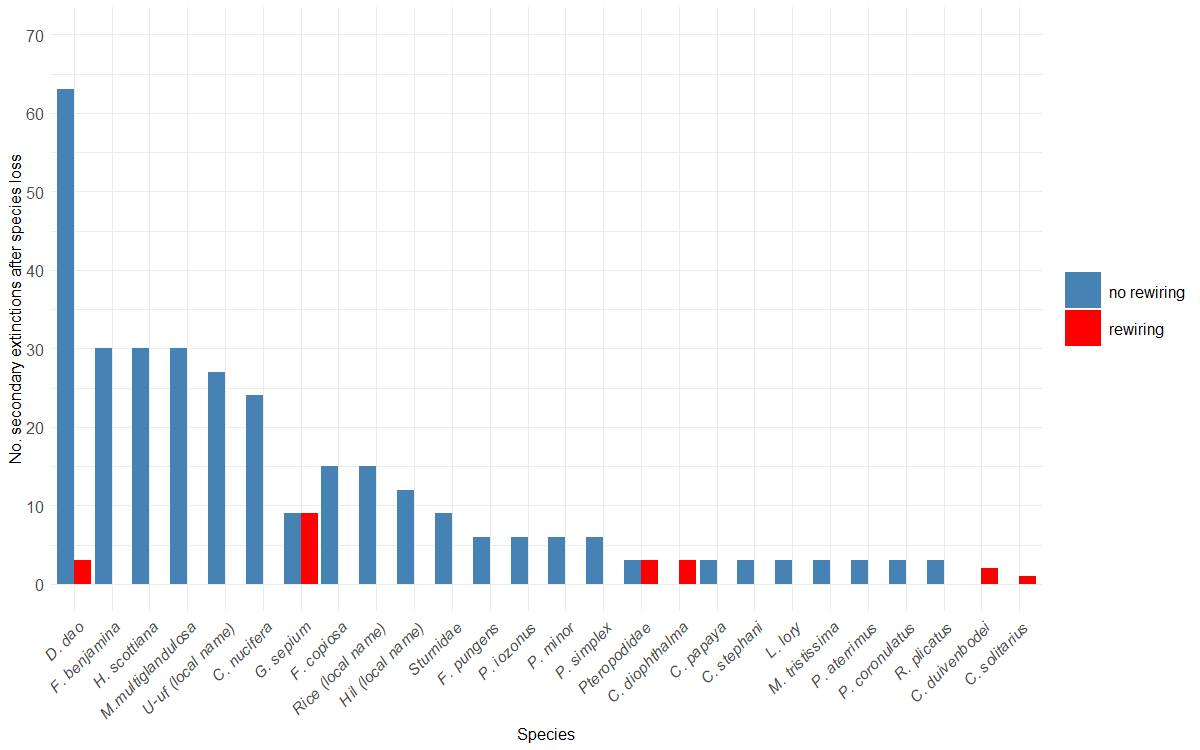

Figure S6: The number of secondary extinctions that occur after specific species are lost from the network following their removal as a primary extinction across all extinction simulations.

Table S3: Relative importance scores of frugivores and plant species for maintaining robust and resilient ecosystem services. Relative importance scores refer to the number of extinction simulations for which a species was considered valuable for maintaining robustness against species loss for the specific ecosystem service. A species is considered valuable when *R* < 0.5 and the species is lost in the first half of primary and secondary extinctions. The following shorthand refers to each ecosystem service category: Fd – Food, Mdcn – Medicine, Ct – Construction and Tools, Fwd – Firewood, Cltr – Culture, Mkt – income from services.; NR = no rewiring; R = rewiring.

|  | Fd |  | Mdcn |  | Ct |  | Fwd |  | Cltr |  | Mkt |  |
| --- | --- | --- | --- | --- | --- | --- | --- | --- | --- | --- | --- | --- |
| Species | NR | R | NR | R | NR | R | NR | R | NR | R | NR | R |
| *Cacatua galerita* | 0 | 0 | 0.133 | 0 | 0.133 | 0 | 0.167 | 0.067 | 0 | 0 | 0 | 0 |
| *Ceyx solitarius* | 0 | 0 | 0.033 | 0 | 0.033 | 0 | 0 | 0 | 0 | 0 | 0 | 0 |
| *Chalcophaps stephani* | 0 | 0 | 0 | 0 | 0 | 0 | 0 | 0.067 | 0 | 0 | 0 | 0 |
| *Chalcopsitta duivenbodei* | 0 | 0 | 0.167 | 0 | 0.233 | 0 | 0.400 | 0 | 0 | 0 | 0 | 0 |
| *Cyclopsitta diophthalma* | 0 | 0 | 0.167 | 0 | 0.167 | 0 | 0.167 | 0 | 0 | 0 | 0 | 0 |
| *Geoffroyus geoffroyi* | 0 | 0 | 0.167 | 0 | 0.233 | 0 | 0.400 | 0 | 0 | 0 | 0 | 0 |
| *Gerygone chrysogaster* | 0 | 0 | 0.033 | 0 | 0.067 | 0 | 0.133 | 0 | 0 | 0 | 0 | 0 |
| *Lorius lory* | 0 | 0 | 0.167 | 0 | 0.133 | 0 | 0.133 | 0 | 0 | 0 | 0 | 0 |
| *Macropygia nigrirostris* | 0 | 0 | 0.033 | 0 | 0.033 | 0 | 0 | 0 | 0 | 0 | 0 | 0 |
| *Mayrimunia tristissima* | 0 | 0 | 0 | 0 | 0 | 0 | 0 | 0 | 0 | 0 | 0 | 0 |
| *Megapodius decollatus* | 0 | 0 | 0.167 | 0 | 0.233 | 0 | 0.400 | 0 | 0 | 0 | 0 | 0 |
| *Micropsitta pusio* | 0 | 0 | 0.167 | 0 | 0.233 | 0 | 0.400 | 0 | 0 | 0 | 0 | 0 |
| *Oriolus szalayi* | 0 | 0 | 0.167 | 0 | 0.233 | 0 | 0.400 | 0 | 0 | 0 | 0 | 0 |
| *Pachycephala simplex* | 0 | 0 | 0 | 0 | 0 | 0 | 0 | 0 | 0 | 0 | 0 | 0 |
| *Paradisaea minor* | 0 | 0 | 0 | 0 | 0 | 0 | 0 | 0 | 0 | 0 | 0 | 0 |
| *Philemon buceroides* | 0 | 0 | 0.167 | 0 | 0.233 | 0 | 0.400 | 0 | 0 | 0 | 0 | 0 |
| *Probosciger aterrimus* | 0 | 0 | 0.167 | 0 | 0.233 | 0 | 0.333 | 0.067 | 0 | 0 | 0 | 0 |
| *Psittaciformes* | 0 | 0 | 0.167 | 0 | 0.233 | 0 | 0.333 | 0.067 | 0 | 0 | 0 | 0 |
| *Pteropodidae* | 0 | 0 | 0.167 | 0 | 0.200 | 0 | 0.233 | 0.066666667 | 0 | 0 | 0 | 0 |
| *Ptilinopus coronulatus* | 0 | 0 | 0 | 0 | 0 | 0 | 0 | 0 | 0 | 0 | 0 | 0 |
| *Ptilinopus iozonus* | 0 | 0 | 0 | 0 | 0 | 0 | 0 | 0.067 | 0 | 0 | 0 | 0 |
| *Ptilinopus nainus* | 0 | 0 | 0.167 | 0 | 0.233 | 0 | 0.400 | 0 | 0 | 0 | 0 | 0 |
| *Ptilinopus pulchellus* | 0 | 0 | 0.167 | 0 | 0.233 | 0 | 0.400 | 0 | 0 | 0 | 0 | 0 |
| *Rhyticeros plicatus* | 0 | 0 | 0.133 | 0 | 0.133 | 0 | 0.167 | 0.067 | 0 | 0 | 0 | 0 |
| *Sturnidae* | 0 | 0 | 0.167 | 0 | 0.167 | 0 | 0.200 | 0.067 | 0 | 0 | 0 | 0 |
| *Abelmoschus manihot* | 0 | 0 | 0 | 0 | 0 | 0 | 0 | 0 | 0 | 0 | 0 | 0 |
| *Amaranthus aupa* | 0 | 0 | 0 | 0 | 0 | 0 | 0 | 0 | 0 | 0 | 0 | 0 |
| *Ananas comosus* | 0 | 0 | 0 | 0 | 0 | 0 | 0 | 0 | 0 | 0 | 0 | 0 |
| *Annona muricata* | 0 | 0 | 0.033 | 0 | 0.033 | 0 | 0 | 0.167 | 0 | 0 | 0 | 0 |
| *Arachis hypogaea* | 0 | 0 | 0.033 | 0 | 0.033 | 0 | 0 | 0.1 | 0 | 0 | 0 | 0 |
| *Areca catechu* | 0 | 0 | 0.167 | 0 | 0.167 | 0 | 0.133 | 0.1 | 0 | 0 | 0 | 0 |
| *Arenga microcarpa* | 0 | 0 | 0.033 | 0 | 0.033 | 0 | 0 | 0.133 | 0 | 0 | 0 | 0 |
| *Arenga spp.* | 0 | 0 | 0.033 | 0 | 0.033 | 0 | 0 | 0.133 | 0 | 0 | 0 | 0 |
| *Artocarpus altilis* | 0 | 0 | 0.167 | 0 | 0.233 | 0 | 0.400 | 0.267 | 0 | 0 | 0 | 0 |
| *Brassica oleracea* | 0 | 0 | 0 | 0 | 0 | 0 | 0 | 0 | 0 | 0 | 0 | 0 |
| *Brassica rapa* | 0 | 0 | 0 | 0 | 0 | 0 | 0 | 0 | 0 | 0 | 0 | 0 |
| *Broussonetia papyrifera* | 0 | 0 | 0 | 0 | 0 | 0 | 0 | 0 | 0 | 0 | 0 | 0 |
| *Calamus longipinna* | 0 | 0 | 0.100 | 0 | 0.133 | 0 | 0.233 | 0.233 | 0 | 0 | 0 | 0 |
| *Canarium indicum* | 0 | 0 | 0.167 | 0 | 0.233 | 0 | 0.400 | 0.267 | 0 | 0 | 0 | 0 |
| *Capsicum frutescens* | 0 | 0 | 0 | 0 | 0 | 0 | 0 | 0 | 0 | 0 | 0 | 0 |
| *Carica papaya* | 0 | 0 | 0.067 | 0 | 0.133 | 0 | 0.300 | 0.067 | 0 | 0 | 0 | 0 |
| *Caryota rumphiana* | 0 | 0 | 0.033 | 0 | 0.033 | 0 | 0 | 0.1 | 0 | 0 | 0 | 0 |
| *Cinnamomum grandifolium* | 0 | 0 | 0.100 | 0 | 0.100 | 0 | 0.067 | 0.133 | 0 | 0 | 0 | 0 |
| *Citrullus lanatus* | 0 | 0 | 0 | 0 | 0 | 0 | 0 | 0 | 0 | 0 | 0 | 0 |
| *Cocos nucifera* | 0 | 0 | 0.033 | 0 | 0.067 | 0 | 0.167 | 0.367 | 0 | 0 | 0 | 0 |
| *Colocasia esculenta* | 0 | 0 | 0 | 0 | 0.067 | 0 | 0.100 | 0.1 | 0 | 0 | 0 | 0 |
| *Cordyline fruiticosa* | 0 | 0 | 0.033 | 0 | 0.050 | 0 | 0.067 | 0.167 | 0 | 0 | 0 | 0 |
| *Cucumis sativas* | 0 | 0 | 0 | 0 | 0 | 0 | 0 | 0 | 0 | 0 | 0 | 0 |
| *Cucumis sp.* | 0 | 0 | 0.100 | 0 | 0.100 | 0 | 0.067 | 0 | 0 | 0 | 0 | 0 |
| *Cucurbita moschata* | 0 | 0 | 0 | 0 | 0 | 0 | 0 | 0 | 0 | 0 | 0 | 0 |
| *Curcurbitaceae* | 0 | 0 | 0 | 0 | 0 | 0 | 0 | 0 | 0 | 0 | 0 | 0 |
| *Cyathea angiensis* | 0 | 0 | 0.067 | 0 | 0.100 | 0 | 0.200 | 0.267 | 0 | 0 | 0 | 0 |
| *Dendrocnide longifolia* | 0 | 0 | 0.100 | 0 | 0.100 | 0 | 0.067 | 0.1 | 0 | 0 | 0 | 0 |
| *Dioscorea alata* | 0 | 0 | 0.033 | 0 | 0.033 | 0 | 0 | 0.133 | 0 | 0 | 0 | 0 |
| *Dioscorea bulbifera* | 0 | 0 | 0 | 0 | 0 | 0 | 0 | 0 | 0 | 0 | 0 | 0 |
| *Dioscorea esculenta* | 0 | 0 | 0 | 0 | 0 | 0 | 0 | 0 | 0 | 0 | 0 | 0 |
| *Dioscorea pentaphylla* | 0 | 0 | 0 | 0 | 0 | 0 | 0.033 | 0.1 | 0 | 0 | 0 | 0 |
| *Dracontomelon dao* | 0 | 0 | 0.167 | 0 | 0.233 | 0 | 0.400 | 0.267 | 0 | 0 | 0 | 0 |
| *Ficus benjamina* | 0 | 0 | 0.167 | 0 | 0.233 | 0 | 0.400 | 0.267 | 0 | 0 | 0 | 0 |
| *Ficus copiosa* | 0 | 0 | 0.167 | 0 | 0.233 | 0 | 0.400 | 0.267 | 0 | 0 | 0 | 0 |
| *Ficus dammaropsis* | 0 | 0 | 0.067 | 0 | 0.133 | 0 | 0.300 | 0.267 | 0 | 0 | 0 | 0 |
| *Ficus pungens* | 0 | 0 | 0.167 | 0 | 0.167 | 0 | 0.133 | 0.1 | 0 | 0 | 0 | 0 |
| *Ficus septica* | 0 | 0 | 0.100 | 0 | 0.100 | 0 | 0.067 | 0.1 | 0 | 0 | 0 | 0 |
| *Ficus virgata* | 0 | 0 | 0.167 | 0 | 0.167 | 0 | 0.200 | 0 | 0 | 0 | 0 | 0 |
| *Geunsia pentandra* | 0 | 0 | 0.167 | 0 | 0.167 | 0 | 0.200 | 0 | 0 | 0 | 0 | 0 |
| *Glycine max* | 0 | 0 | 0.033 | 0 | 0.033 | 0 | 0 | 0.1 | 0 | 0 | 0 | 0 |
| *Gnetum gnemon* | 0 | 0 | 0.100 | 0 | 0.167 | 0 | 0.333 | 0.1 | 0 | 0 | 0 | 0 |
| *Homalanthus novoguineensis* | 0 | 0 | 0.033 | 0 | 0.033 | 0 | 0 | 0.267 | 0 | 0 | 0 | 0 |
| *Hornstedtia scottiana* | 0 | 0 | 0.100 | 0 | 0.100 | 0 | 0.067 | 0.167 | 0 | 0 | 0 | 0 |
| *Hydriastele costata* | 0 | 0 | 0.133 | 0 | 0.133 | 0 | 0.133 | 0.167 | 0 | 0 | 0 | 0 |
| *Inocarpus fagifer* | 0 | 0 | 0.033 | 0 | 0.033 | 0 | 0.100 | 0 | 0 | 0 | 0 | 0 |
| *Instia bijuga* | 0 | 0 | 0.167 | 0 | 0.167 | 0 | 0.133 | 0.267 | 0 | 0 | 0 | 0 |
| *Ipomoea batatas* | 0 | 0 | 0 | 0 | 0 | 0 | 0 | 0.1 | 0 | 0 | 0 | 0 |
| *Laportea decumana* | 0 | 0 | 0.100 | 0 | 0.100 | 0 | 0.067 | 0 | 0 | 0 | 0 | 0 |
| *Leucaena leucocephala* | 0 | 0 | 0 | 0 | 0 | 0 | 0 | 0.1 | 0 | 0 | 0 | 0 |
| *Litsea* | 0 | 0 | 0 | 0 | 0 | 0 | 0 | 0 | 0 | 0 | 0 | 0 |
| *Lycopersicon esculentum* | 0 | 0 | 0 | 0 | 0 | 0 | 0 | 0 | 0 | 0 | 0 | 0 |
| *Macaranga aleuritoides* | 0 | 0 | 0.033 | 0 | 0.033 | 0 | 0 | 0 | 0 | 0 | 0 | 0 |
| *Mangifera indica* | 0 | 0 | 0.167 | 0 | 0.233 | 0 | 0.400 | 0.1 | 0 | 0 | 0 | 0 |
| *Mangifera minor* | 0 | 0 | 0.167 | 0 | 0.233 | 0 | 0.400 | 0.267 | 0 | 0 | 0 | 0 |
| *Manihot esculenta* | 0 | 0 | 0 | 0 | 0 | 0 | 0 | 0.267 | 0 | 0 | 0 | 0 |
| *Melanolepis multiglandulosa* | 0 | 0 | 0.167 | 0 | 0.233 | 0 | 0.400 | 0 | 0 | 0 | 0 | 0 |
| *Metroxylon sagu* | 0 | 0 | 0.033 | 0 | 0.033 | 0 | 0.067 | 0.267 | 0 | 0 | 0 | 0 |
| *Moraceae* | 0 | 0 | 0.167 | 0 | 0.167 | 0 | 0.167 | 0.267 | 0 | 0 | 0 | 0 |
| *Musa peekelii* | 0 | 0 | 0.133 | 0 | 0.167 | 0 | 0.200 | 0.1 | 0 | 0 | 0 | 0 |
| *Musa sp.* | 0 | 0 | 0.067 | 0 | 0.133 | 0 | 0.167 | 0 | 0 | 0 | 0 | 0 |
| *NARI taro* | 0 | 0 | 0 | 0 | 0 | 0 | 0 | 0.1 | 0 | 0 | 0 | 0 |
| *Nasturtium officinale* | 0 | 0 | 0 | 0 | 0 | 0 | 0 | 0 | 0 | 0 | 0 | 0 |
| *Nastus elatus* | 0 | 0 | 0.167 | 0 | 0.233 | 0 | 0.367 | 0 | 0 | 0 | 0 | 0 |
| *Oryza sativa* | 0 | 0 | 0 | 0 | 0 | 0 | 0 | 0.367 | 0 | 0 | 0 | 0 |
| *Pandanus conoideus* | 0 | 0 | 0.033 | 0 | 0.033 | 0 | 0 | 0 | 0 | 0 | 0 | 0 |
| *Pangium edule* | 0 | 0 | 0.167 | 0 | 0.233 | 0 | 0.400 | 0.133 | 0 | 0 | 0 | 0 |
| *Passiflora edulis* | 0 | 0 | 0.033 | 0 | 0.033 | 0 | 0 | 0.267 | 0 | 0 | 0 | 0 |
| *Phaseolus vulgaris* | 0 | 0 | 0 | 0 | 0 | 0 | 0 | 0.1 | 0 | 0 | 0 | 0 |
| *Piper betel* | 0 | 0 | 0.100 | 0 | 0.100 | 0 | 0.067 | 0 | 0 | 0 | 0 | 0 |
| *Pipturus argenteus* | 0 | 0 | 0 | 0 | 0 | 0 | 0 | 0.167 | 0 | 0 | 0 | 0 |
| *Planchonella maclayana* | 0 | 0 | 0.100 | 0 | 0.167 | 0 | 0.333 | 0 | 0 | 0 | 0 | 0 |
| *Pometia Pinnata* | 0 | 0 | 0.167 | 0 | 0.233 | 0 | 0.400 | 0.267 | 0 | 0 | 0 | 0 |
| *Psidium guajava* | 0 | 0 | 0.100 | 0 | 0.133 | 0 | 0.233 | 0.267 | 0 | 0 | 0 | 0 |
| *Psophocarpus tetragonolobus* | 0 | 0 | 0 | 0 | 0 | 0 | 0 | 0.267 | 0 | 0 | 0 | 0 |
| *Pterocarpus indicus* | 0 | 0 | 0.100 | 0 | 0.100 | 0 | 0.067 | 0 | 0 | 0 | 0 | 0 |
| *Saccharum edule* | 0 | 0 | 0 | 0 | 0 | 0 | 0 | 0.1 | 0 | 0 | 0 | 0 |
| *Saccharum officinarum* | 0 | 0 | 0 | 0 | 0 | 0 | 0 | 0 | 0 | 0 | 0 | 0 |
| *Solanum melongena* | 0 | 0 | 0 | 0 | 0 | 0 | 0 | 0 | 0 | 0 | 0 | 0 |
| *Spathodea campanulata* | 0 | 0 | 0.133 | 0 | 0.133 | 0 | 0.133 | 0 | 0 | 0 | 0 | 0 |
| *Syzygium amplum* | 0 | 0 | 0.067 | 0 | 0.133 | 0 | 0.300 | 0 | 0 | 0 | 0 | 0 |
| *Tecomanthe dendrophila* | 0 | 0 | 0.033 | 0 | 0.033 | 0 | 0 | 0.267 | 0 | 0 | 0 | 0 |
| *Terminalia catappa* | 0 | 0 | 0.167 | 0 | 0.233 | 0 | 0.400 | 0.1 | 0 | 0 | 0 | 0 |
| *Terminalia kaernbachii* | 0 | 0 | 0.167 | 0 | 0.233 | 0 | 0.333 | 0.267 | 0 | 0 | 0 | 0 |
| *Theobroma cacao* | 0 | 0 | 0 | 0 | 0 | 0 | 0 | 0 | 0 | 0 | 0 | 0 |
| *Vigna unguiculata* | 0 | 0 | 0 | 0 | 0 | 0 | 0 | 0 | 0 | 0 | 0 | 0 |
| *Xanthosoma sagittifolium* | 0 | 0 | 0 | 0 | 0 | 0 | 0 | 0 | 0 | 0 | 0 | 0 |
| *Zea mays* | 0 | 0 | 0 | 0 | 0 | 0 | 0 | 0 | 0 | 0 | 0 | 0 |
| *Zingiber officinale* | 0 | 0 | 0.033 | 0 | 0.067 | 0 | 0.167 | 0 | 0 | 0 | 0 | 0 |
